# Sustained Glucose Turnover Flux Distinguishes Cancer Cachexia from Nutrient Limitation

**DOI:** 10.1101/2025.05.15.654370

**Authors:** Young-Yon Kwon, Yanshan Liang, Maria Gomez-Jenkins, Mujmmail Ahmed, Guangru Jiang, Juliya Hsiang, David Y. Lewis, Tobias Janowitz, Marcus D. Goncalves, Eileen White, Sheng Hui

## Abstract

Cancer cachexia is an involuntary weight loss condition characterized by systemic metabolic disorder. A comprehensive flux characterization of this condition however is lacking. Here, we systematically isotope traced eight major circulating nutrients in mice bearing cachectic C26 tumors (cxC26) and food intake-matched mice bearing non-cachectic C26 tumors (ncxC26). We found no difference in whole-body lipolysis and proteolysis, ketogenesis, or fatty acid and ketone oxidation by tissues between the two groups. In contrast, compared to ncxC26 mice ad libitum, glucose turnover flux decreased in food intake-controlled ncxC26 mice but not in cxC26 mice. Similarly, sustained glucose turnover flux was observed in two autochthonous cancer cachexia models despite reduced food intake. We identified glutamine and alanine as responsible for sustained glucose production and tissues with altered use of glucose and lactate in cxC26 mice. We provide a comprehensive view of metabolic alterations in cancer cachexia revealing those distinct from decreased nutrient intake.

**Highlights:**

- Quantitative fluxomics of cancer cachexia under matched food intake and body weight
- Intact lipolysis, proteolysis, ketogenesis, and lipid oxidation in cachectic mice
- Sustained glucose consumption in cachectic mice despite reduced food intake
- Increased glucose production from glutamine and alanine in cachectic mice

## Introduction

Cancer cachexia is a tumor-induced debilitating condition characterized by involuntary body weight loss and tissue wasting (Baracos et al., 2018; Evans et al., 2008; Ferrer et al., 2023). It affects up to 80% of advanced cancer patients and is associated with poor quality of life and worse survival. Progressive tissue wasting during cachexia is thought to be driven, in part, by excessive lipolysis in adipose tissues, leading to the rapid mobilization of triglycerides and depletion of fat stores, and by increased skeletal muscle proteolysis, which results in the breakdown of structural and contractile proteins. In addition to wasting of adipose tissues and muscles, a range of metabolic alterations have been reported in cancer cachexia, including abnormal thermogenesis in adipose tissues, reduced mitochondrial metabolism in muscles, and perturbed systemic metabolism with changes to the Cori cycle, plasma lipid profile, insulin sensitivity, and feeding behavior. Thus, cancer cachexia is viewed as a complex metabolic disorder with progressive tissue wasting (Berriel Diaz et al., 2024; Evans et al., 2008; Ferrer et al., 2023; Tisdale, 2009)

To understand the mechanisms underlying this metabolic remodeling, prior investigations have examined systemic metabolism in cancer cachexia using gene expression analysis and metabolomic profiling. For example, based on elevated expression of lipolysis enzymes, particularly hormone-sensitive lipase (HSL), increased lipolysis in adipose tissues has been suggested as a cause of adipose tissue wasting in cancer cachexia (Agustsson et al., 2007; Cao et al., 2010). Similarly, elevated expression levels of E3 ubiquitin ligases, MuRF1 and Atrogin-1, are used as biomarkers for increased protein degradation and consequently skeletal muscle atrophy (Liang et al., 2025; Sukari et al., 2016). However, the functional in vivo activity of these enzymes—specifically their fluxes within intact physiological systems— remains largely unquantified, preventing definitive conclusions regarding these processes’ role in tissue wasting. In addition to gene expression data, some studies have tried to infer metabolic changes in cancer cachexia from steady state metabolite levels using metabolomics on samples from cancer cachexia patients and preclinical models (Cala et al., 2018; Der-Torossian et al., 2013; Goncalves et al., 2018; Pin et al., 2019; Potgens et al., 2021). These studies have shown alterations in the levels of metabolites related to glycolysis, tricarboxylic acid cycle (TCA cycle), and branch-chain amino acid metabolism. While steady state levels of metabolites provide valuable metabolic fingerprints of cancer cachexia, they do not capture the dynamic nature of metabolism.

In vivo flux analysis allows determination of dynamic aspects of metabolism and insights on the production, utilization, and interconversion of metabolites in intact animals or humans (Faubert and DeBerardinis, 2017; Robert R. Wolfe, 2004). A small number of studies pioneered flux characterization in patients with cancer cachexia. Two papers reported higher glucose turnover flux in cancer patients with progressive weight loss compared to healthy volunteers (Holroyde et al., 1984) or cancer patients with stable weight (Holroyde et al., 1975). Regarding lipolysis flux, Jeevanandam et al. found no change in cachectic patients compared to normal subjects by measuring glycerol turnover flux (Jeevanandam et al., 1986). In contrast, Legaspi et al. measured higher glycerol and free fatty acid (FFA) turnover fluxes in cancer patients with weight loss than reported values for healthy subjects (Legaspi et al., 1987). Proteolysis flux has also been measured. Norton et al. showed an increased whole-body protein turnover in malnourished cancer patients (Norton et al., 1981), while Lundholm et al. demonstrated stable protein degradation in cancer patients with weight loss (Lundholm et al., 1982). These studies suggest that while glucose turnover flux is consistently elevated in cancer patients with weight loss, findings on lipolysis and proteolysis fluxes are variable, highlighting the heterogeneity of metabolic alterations in cachexia and the need for further flux studies to clarify tissue-specific mechanisms.

Metabolic measurements including flux studies can be confounded by multiple factors that are present in cachexia. One important metabolic confounding factor is food intake. Reduced food intake, or anorexia, is a major symptom in patients with cancer cachexia (Ezeoke and Morley, 2015), and also in many preclinical cancer cachexia models (Flint et al., 2016; Kim-Muller et al., 2023; Liang et al., 2025; Queiroz et al., 2022). Reduced food intake induces significant metabolic alterations in the body, independent of diseases (Casanova et al., 2019; Collet et al., 2017; Garcia-Flores et al., 2021; Xie et al., 2022). For instance, dietary restriction alone can lead to loss of lean and fat mass, lower fasting glucose levels, and reduced energy expenditure (Di Francesco et al., 2024; Redman et al., 2018). Therefore, it is critical to separate the effects of reduced food intake while studying metabolic alterations in cancer cachexia (Emery, 1999; Liang et al., 2025). Other confounding factors of metabolic measurements include body weight and composition, which can potentially influence metabolite levels and fluxes. While body weight’s influence can be disentangled using statistical analysis (Speakman, 2013), in general it is unclear how body weight and composition affect fluxes through different metabolic pathways. Thus, flux alterations measured between cachectic and healthy subjects could be due to their different body weight and composition and may not reflect dysregulation of metabolism in cancer cachexia. Therefore, to reveal metabolic alterations that are inherent to cachexia it is crucial to control for these metabolic confounders while measuring metabolism.

Here, with mass spectrometry-based in vivo isotope tracing, we systemically determined fluxes of eight major circulating nutrients in the colon carcinoma 26 (C26) model of cancer cachexia. As shown in a separate study by us (Liang et al., 2025), while food intake was matched, cachectic C26 mice and their non-cachectic control group exhibited almost identical energy expenditure, body weight loss, and body composition, presenting a highly controlled condition for revealing “cachexia-inherent” metabolic alterations that cannot simply be explained by those confounding factors of metabolism. In this rigorously controlled experimental system, we found no change in whole-body lipolysis, whole-body proteolysis, or ketogenesis flux in cachectic animals, contrary to current views in literature. In contrast, we found changes in the metabolism of glucose and related nutrients including glutamine, alanine, and lactate. Specifically, we found sustained glucose production and utilization despite reduced food intake in the C26 model and two separate genetically modified mouse models of lung cancer cachexia. We further identified the nutrient sources and tissue sinks responsible for the changes in glucose turnover flux in cachectic mice, as well as tissues with altered fuel selection. Our flux results provide a comprehensive view of food intake- and body weight-independent energy metabolism in cancer cachexia, revealing a glucose-centric remodeling of metabolism.

## Results

### A highly controlled experimental system for probing cachexia-inherent metabolic alterations in cancer cachexia

We aimed to systemically evaluate alteration of metabolic fluxes in cancer cachexia using the C26 cancer cachexia mouse model, which displays robust cachectic phenotypes (Bonetto et al., 2016). Mice bearing cachectic C26 tumors (cxC26) show more than 15% of body weight loss within two weeks after cancer cell injection while mice bearing the non-cachectic C26 tumors (ncxC26) maintained body weight with similar tumor growth (Kwon and Hui, 2024) (Figure 1A). An important symptom of cancer cachexia is reduced food intake, or anorexia. This phenotype is captured in the C26 model. While ad libitum, cxC26 mice but not ncxC26 mice exhibit decreasing food intake from day 8 post cancer cell injection (Figure S1), mirroring their body weight curves. Food intake is a major determinant of metabolism and thus anorexia can cause changes in metabolic fluxes. To reveal metabolic changes that cannot simply be accounted for by reduced food intake, here in our flux studies we used a control group that had equal food intake as the cachectic group. Specifically, we implemented an isocaloric (ICa) feeding strategy by gradually reducing the food provided to single housed cxC26 and ncxC26 mice from day 8 to 12 (Figure 1B). The exact amount provided daily was based on the food intake measured for cxC26 mice ad libitum (day 8: 3.1g, day 9: 2.8g, day 10: 1.7g, and day 11: 1.1g) as shown in Figure S1. Moreover, the daily food amount was divided into 3 equal portions that were dispensed every 4 hours during the nighttime to account for any changes related to long fasting intervals (Pak et al., 2021). To ensure equal food intake, we excluded cxC26 mice that did not finish the provided food from our experiments.

**Figure 1.**
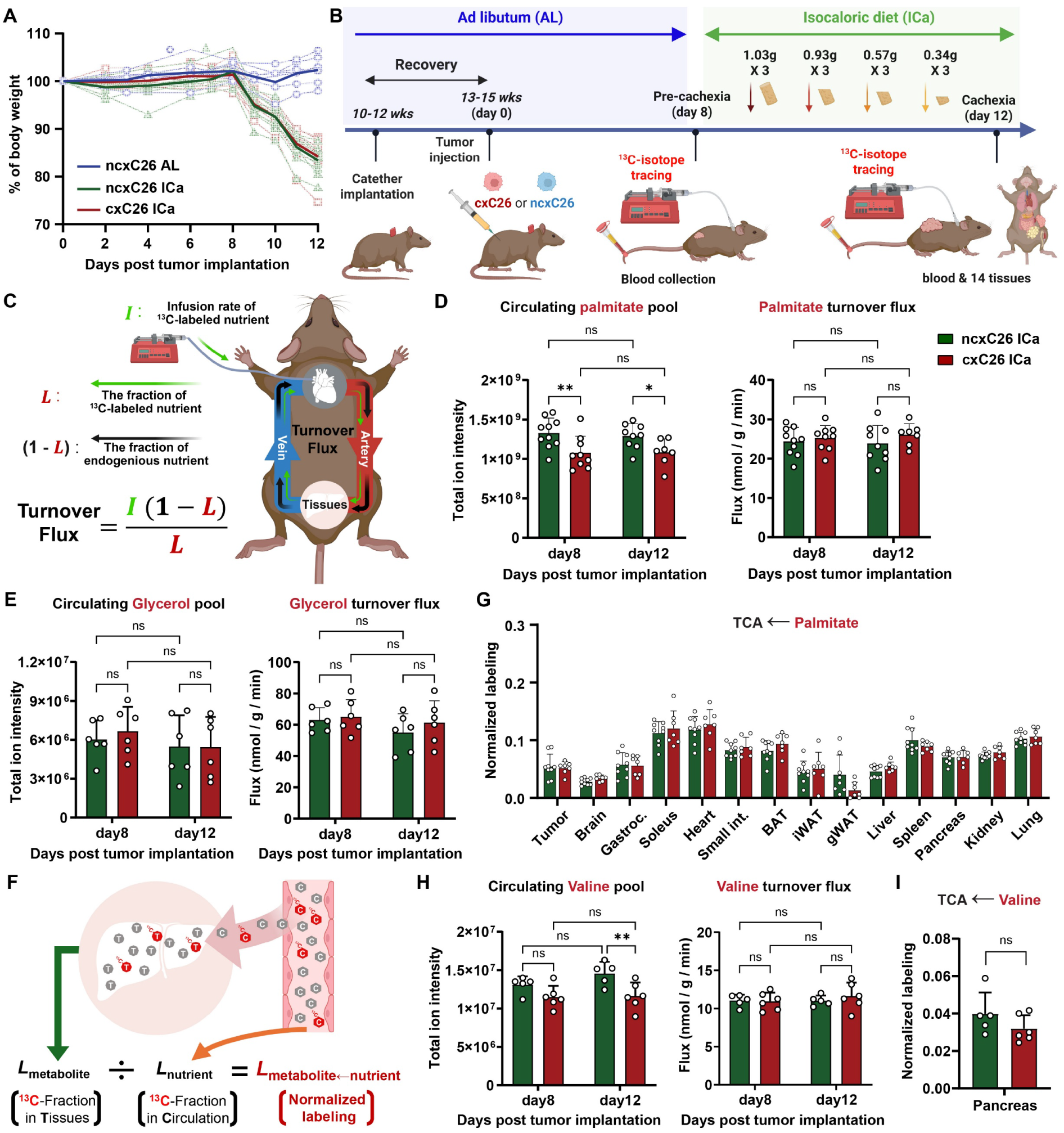
Whole-body lipolysis and proteolysis were not elevated in cachectic mice. (A) Daily body weight of cxC26 and ncxC26 mice under isocaloric feeding (ICa) or ad libitum (AL). (n= 7-10) (B) Illustration of experimental design for isotope tracing. cxC26 and ncxC26 mice were fed equal amount of food from day 8 to day 12 post tumor implantation (Isocaloric feeding; ICa), based on the daily food intake of cxC26 mice ad libitum (see Figure S1b). Isotope tracer infusion was performed at pre-cachectic (day 8) and cachectic (day 12) stages. (C) Illustration of the concept of turnover flux and its calculation from tracer infusion rate (I) and isotope-labeled fraction of traced nutrient in the circulation (L). (D) Circulating palmitate pool and turnover flux in cxC26 and ncxC26 mice under isocaloric feeding at day 8 and day 12. (n= 7-10) (E) Circulating glycerol pool and turnover flux in cxC26 and ncxC26 mice under isocaloric feeding at day 8 and day 12. (n= 6) (F) Illustration of using normalized labeling of TCA intermediates such as malate by a circulating nutrient as the nutrient’s contribution to TCA cycle. (G) Normalized labeling of malate in different tissues under ^13^C-palmitate infusion at day 12. (n= 7-9) (H) Circulating valine pool and turnover flux in cxC26 and ncxC26 mice under isocaloric feeding at day 8 and day 12. (n= 5-6) (I) Normalized labeling of malate in pancreas under ^13^C-valine infusion at day 12. (n= 5-6) Data are shown as mean ± s.d. Significance of the differences: (D,E, H and I) ns: non-significance, *P < 0.05, ** P < 0.01, *** P < 0.001 between groups by two-way ANOVA or (G) no symbol: not significant, *FDR < 0.05, ** FDR < 0.01, *** FDR < 0.001 between groups by two-tailed t-test with multiple corrections.

As shown by us in a separate study, with this isocaloric feeding strategy, cxC26 and ncxC26 mice are indistinguishable in many physiological parameters which are potential confounding factors of metabolism, including energy expenditure, energy excretion, body weight, fat mass, and lean mass (Liang et al., 2025). The similar body weight loss between the two groups was confirmed in this study (Figure 1A). Accordingly, this experimental paradigm offers an exquisitely controlled framework ideally suited for delineating metabolic perturbations intrinsic to cachexia. Note that compared to the food intake-matched ncxC26 mice, cxC26 mice still exhibit impaired physical performance, a key phenotype of cancer cachexia (Liang et al., 2025).

To quantify fluxes, we employed continuous infusion of isotope tracers to these mice. To preserve physiological fidelity, we infused conscious and free-moving mice through a catheter implanted at the jugular vein. For a systematic examination of energy metabolism, we infused separately eight uniformly ^13^C-labeled tracers (palmitate, glycerol, 3-hydroxybutyrate, glucose, lactate, glutamine, alanine and valine) representing all major circulating nutrients (Hui et al., 2020; Yuan et al., 2025) at both pre-cachectic (day 8; before body weight loss and anorexia) and cachectic stage (day 12) (Figure 1B). Thus, we have established an experimental setup for systematically analyzing energy metabolic flux changes that are inherent to cachexia.

### No alterations in whole-body lipolysis flux or tissue fatty acid oxidation in cachectic mice

Loss of fat is a common phenotype in cancer cachexia, observed in patients and animal models, including cxC26 mice (Kwon and Hui, 2024; Liang et al., 2025). Though elevated expression levels of adipose lipase genes such as *Atgl* and *Hsl* have been reported in cancer cachexia (Agustsson et al., 2007; Kir et al., 2014), direct assessment of lipolysis flux in vivo has rarely been conducted. Lipolysis refers to the process of tissue triglycerides being broken down and released as fatty acids and glycerol in the circulation. Thus, the turnover flux of either a circulating fatty acid or glycerol can be taken as lipolysis flux at the whole-body level. To quantify turnover flux of a fatty acid, we infused uniformly ^13^C-labeled palmitate into cxC26 and ncxC26 mice at pre-cachectic and cachectic stage under isocaloric feeding. The fraction of ^13^C-labeled palmitate in the blood circulation when it reached steady state was measured using liquid chromatography-mass spectrometry (LC-MS) to calculate palmitate turnover flux (Hui et al., 2017) (Figure 1C). Notwithstanding perturbations in the circulating palmitate pool, palmitate turnover flux remained unaltered in cxC26 mice at day 12 compared to either day 8 or ncxC26 mice at day 12 (Figure 1D). To validate this result, we also infused the other product of lipolysis, glycerol. Consistent with palmitate turnover flux, glycerol turnover flux in cxC26 mice at day 12 was not altered compared to either day 8 or ncxC26 mice at day 12. The glycerol pool also remained unchanged across all conditions (Figure 1E). Collectively, these data demonstrate that systemic lipolysis flux remained unaltered in cachectic mice, challenging longstanding assumptions regarding adipose catabolism in this context.

Although there was no detectable difference in systemic lipolysis flux, we sought to examine the tissue-level utilization of fatty acids. As fatty acids are a major fuel for tissues, we evaluated the contribution of circulating fatty acids to the TCA cycle, the predominant energy production pathway, in different tissues. For this, we measured the ^13^C-labeling of the TCA intermediate malate in different tissues of mice under ^13^C-palmitate infusion and divided it by the labeling of circulating palmitate (Figure 1F). This normalized labeling of malate in a tissue is taken as the contribution from circulating palmitate to that tissue’s TCA cycle. There was no significant change of palmitate contribution in any of the tissues between cachectic and non-cachectic mice under isocaloric feeding (Figure 1G), indicating intact fatty oxidation in cachectic mice. Moreover, comparison of labeling of all detectable tissue metabolites showed no significant changes in any of the measured tissues in cachectic mice (Figure S2A and B), supporting intact utilization of fatty acids in cachectic mice. Altogether, in the C26 model, there was no change in whole body lipolysis or fatty acid utilization by tissues.

### No alterations in whole-body proteolysis flux or essential amino acid oxidation by tissues in cachectic mice

Muscle wasting is another primary phenotype of cancer cachexia, which is captured by the cxC26 mice (Kwon and Hui, 2024). Elevated degradation of proteins, or proteolysis, is often cited as a cause of reduced muscle mass. To investigate this, we aimed to quantify the proteolysis flux. We infused uniformly ^13^C-labeled valine to determine its turnover flux. As an essential amino acid, the sole source for circulating valine in fasted mice is tissue proteins. Thus, valine turnover flux in fasted mice reflects the whole-body proteolysis. Unexpectedly, valine turnover flux was not changed in cxC26 mice compared to ncxC26 mice, despite reduced circulating valine pool in cxC26 mice at cachectic stage (Figure 1H). Moreover, valine’s contribution to tissue metabolites including TCA cycle intermediates showed no change between the groups (Figure 1I and Figure S3). Though it remains a possibility that proteolysis can change in individual tissues such as muscle in cachexia, the results indicate that at the whole-body level proteolysis was not elevated in the C26 model.

### No alterations in ketogenesis or tissue ketone utilization in cachectic mice

An important energy nutrient that is closely related to fatty acid metabolism is ketone bodies, which are primarily produced from fatty acids (Newman and Verdin, 2014). Previous studies have implicated altered production of ketone bodies, or ketogenesis, in cancer cachexia (Flint et al., 2016; Goncalves et al., 2018). To determine whether ketone metabolism is altered in our C26 model, we infused uniformly ^13^C-labeled 3-hydroxybutyrate (3-HB) to cxC26 and ncxC26 mice under isocaloric feeding. Compared to day 8, the pool of circulating 3-HB was significantly elevated in ncxC26 mice at day 12, which is normal response to reduced food intake. Interestingly, the extent of elevation was the same in the cxC26 mice (Figure 2A). This similarity was also observed in the turnover flux of 3-HB, which increased to similar levels in both groups from day 8 to day 12 (Figure 2A). These results point to normal response of ketogenesis to reduced food intake in cachectic mice.

**Figure 2.**
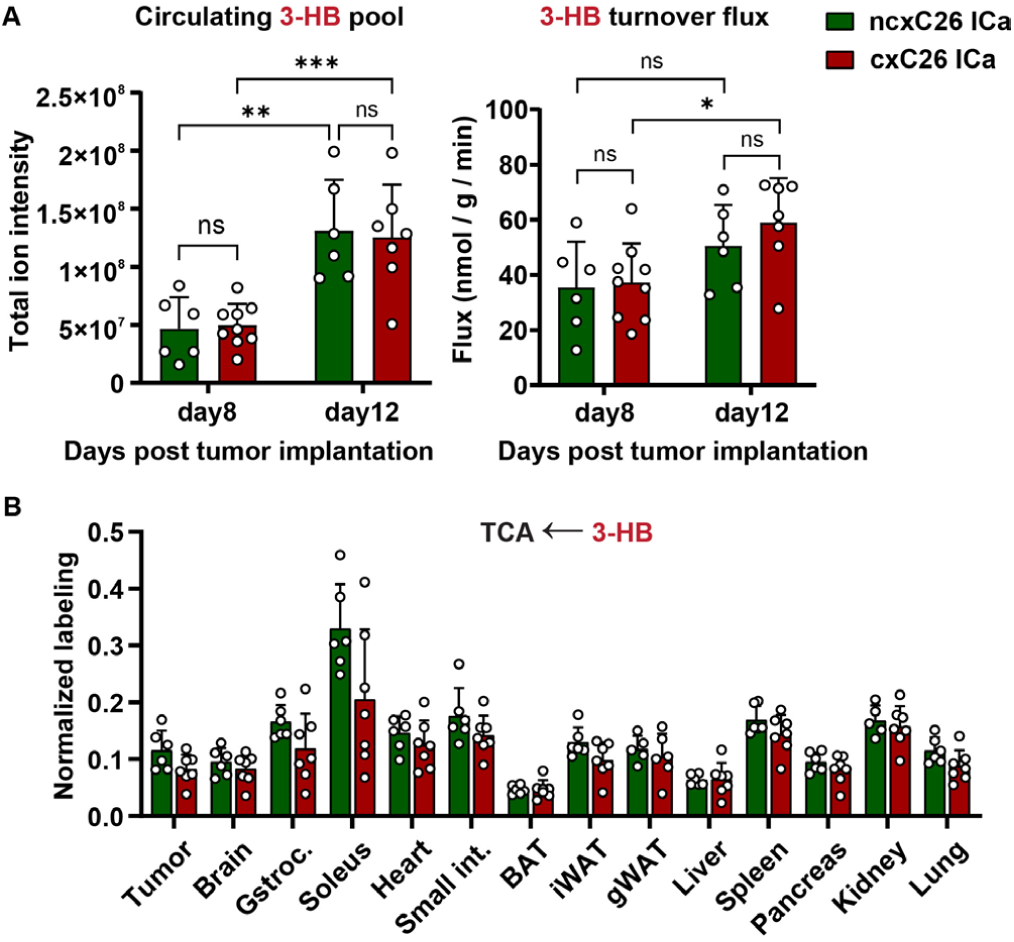
Ketogenesis was elevated in cachectic mice, but only due to reduced food intake. (A) Circulating 3-hydroxybutyrate (3-HB) pool and turnover flux in cxC26 and ncxC26 mice under isocaloric feeding (ICa) at day 8 and day 12 post tumor implantation. (n= 6-9) (B) Normalized labeling of malate in tissues under ^13^C-3-HB infusion at day 12. (n= 6-7) Data are shown as mean ± s.d. Significance of the differences: (A) ns: non-significance, *P < 0.05, ** P < 0.01, *** P < 0.001 between groups by two-way ANOVA or (B) no symbol: not significant, *FDR < 0.05, ** FDR < 0.01, *** FDR < 0.001 between groups by two-tailed t-test with multiple corrections.

We next investigated utilization of 3-HB by tissues. We first examined contribution of 3-HB as a fuel to tissues and found no significant change in any of the tissues measured in cachectic mice (figure 2B). We then analyzed contribution of 3-HB to all metabolites with measurable labeling. Although only two tissue metabolites (GMP in the brain and alanine in the tumor) of 3-HB showed significant labeling changes, PCA analysis revealed distinct labeling profiles in the tumor, kidney, and pancreas of cxC26 mice (Figure S4). These results suggest some small overall trend in the labeling of tissue metabolites rather than significant changes in select metabolites. Altogether, we conclude that ketone metabolism is nearly normal in cachectic mice.

### Sustained glucose turnover flux in cachectic mice despite reduced food intake

To continue probing energy metabolism in cachexia, we next moved to glucose to determine its production and utilization fluxes. We infused uniformly ^13^C-labeled glucose into cxC26 and ncxC26 mice under isocaloric feeding. As shown in Figure 3A, both groups had decreased levels of circulating glucose at day 12 compared to day 8, with slightly lower levels in the cxC26 group. This reduction in the circulating glucose levels was expected due to the restriction of food intake. However, we identified a clear difference between the groups in glucose turnover fluxes. While the ncxC26 group decreased their glucose turnover flux from day 8 to day 12, the cxC26 group surprisingly maintained their glucose turnover flux. It was possible that the lower glucose turnover flux in ncxC26 mice at day 12 was due to the presence of ncxC26 tumors instead of food restriction. To test this possibility, we also infused ^13^C-labeled glucose into ncxC26 mice ad libitum at day 12 and found no change in glucose turnover flux compared to day 8 (Figure 3A), demonstrating that the glucose flux reduction in the isocalorically fed ncxC26 mice was due to food restriction. These results reveal that despite a reduction in circulating glucose levels, cachectic mice fail to downregulate glucose turnover flux in response to reduced food intake, indicating an impaired regulatory adaptation of glucose metabolism in cachexia.

**Figure 3:**
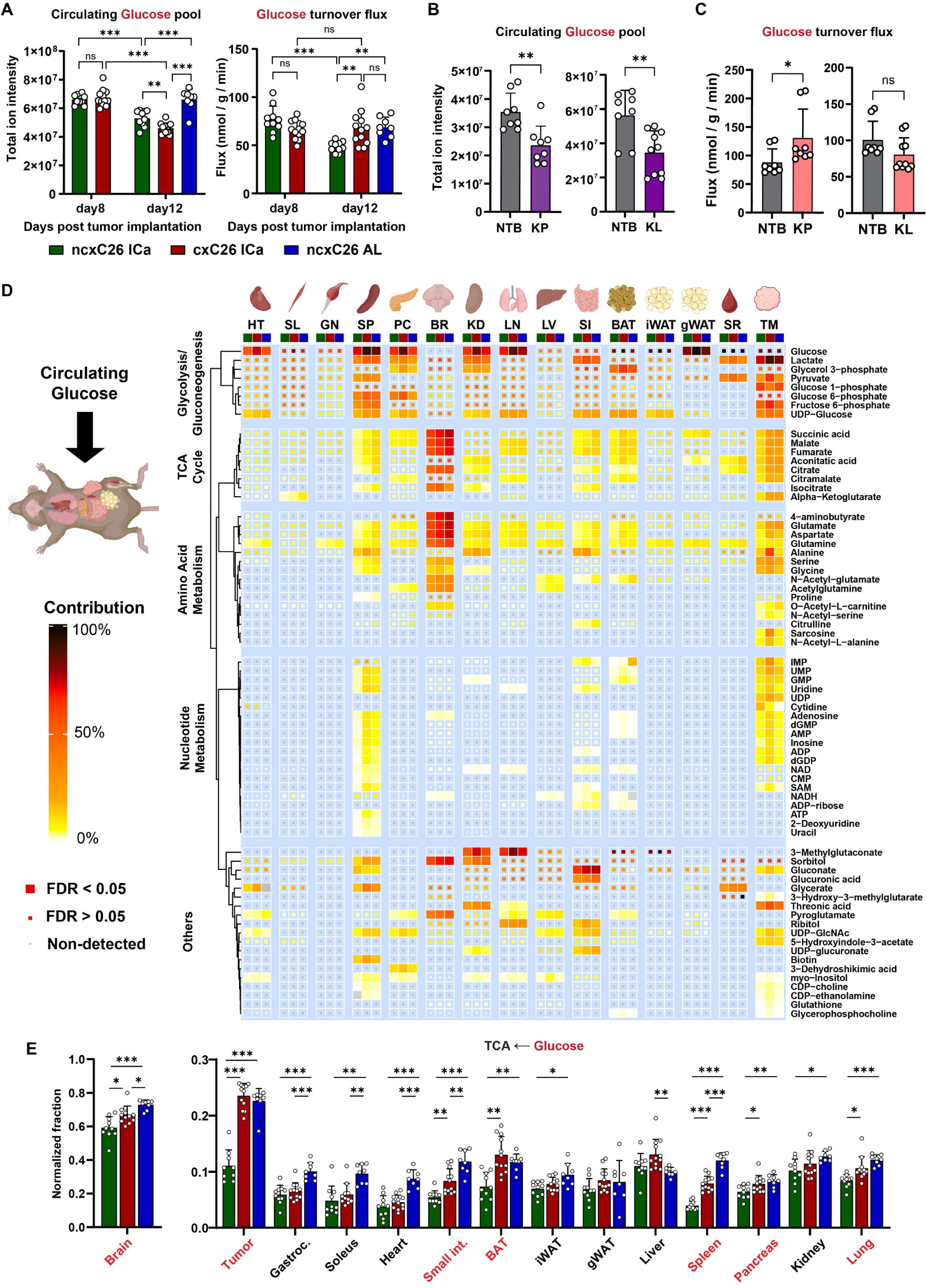
Glucose turnover flux was sustained in cachectic mice while it was decreased in food intake-controlled non-cachectic mice. (A) Circulating glucose pool and turnover flux in cxC26 and ncxC26 mice under isocaloric feeding (ICa), and ncxC26 mice ad libitum (AL), at day 8 and day 12 post tumor implantation. (n= 8-12) (B, C) Circulating glucose pool and turnover flux in cachectic (B) KP and (C) KL mice compared to their respective non-tumor-bearing control mice (NTB). (n= 8-10) (D) Heatmap showing normalized labeling of significantly changed metabolites in tissues and tumors under ^13^C-glucose infusion at day 12. Metabolites were clustered by metabolic pathways. Color indicates magnitude of normalized labeling (or contribution from glucose). A significantly changed metabolite was defined by an FDR < 0.05 in at least one tissue between cxC26 ICa and ncxC26 ICa, with a normalized labeling greater than 1% in that tissue. Filled entries for a metabolite in a tissue indicate significant change of the metabolite’s labeling across the three mouse groups in that tissue. (n= 8-11) (E) Normalized labeling of malate in tissues and tumors under ^13^C-glucose infusion at day 12. Tissues with significant differences between cxC26 ICa and ncxC26 ICa are highlighted in red. (n= 8-11) Data are shown as mean ± s.d. Significance of the differences: (A-C) ns: non-significance, *P < 0.05, ** P < 0.01, *** P < 0.001 between groups by two-way ANOVA or (D, E) no symbol: not significant, *FDR < 0.05, ** FDR < 0.01, *** FDR < 0.001 between groups by two-tailed t-test with multiple corrections.

We next sought to determine if our findings regarding glucose metabolism are applicable to other cancer cachexia models. For this, we measured glucose turnover flux in two different non-small cell lung cancer cachexia models, KP (*Kras^G12D/+^;p53^-/-^*) (Jackson et al., 2005) and KL (*Kras^G12D/+^;Lkb1^-/-^*) (Ji et al., 2007). Both KP (Figure S5) (Verlande et al., 2022) and KL (Queiroz et al., 2022) models have reductions in food intake over time and develop cachexia. Similar to our studies in the C26 model, compared to their respective non-tumor-bearing control mice ad libitum, circulating glucose pools were reduced in both KP and KL mice ad libitum (Figure 3B and C). Despite this reduction, glucose turnover flux was significantly higher in KP mice and maintained in KL mice (Figure 3B and C). These results mirror the finding in the C26 model and suggest that this lack of downregulation of glucose turnover flux in response to reduced food intake is a conserved feature of cachexia.

### Sustained glucose utilization by specific tissues of cachectic mice

The sustained glucose turnover flux in cxC26 mice points to altered glucose utilization by tissues in these mice. We thus sought to evaluate tissue glucose utilization in these mice. At the end of ^13^C glucose infusion, we measured labeling of metabolites in 13 major tissues as well as tumor. As shown in Figure 3D, there were widespread changes in contribution of glucose to tissue metabolites (defined as tissue metabolite labeling divided by circulating glucose labeling, or normalized labeling) between the isocalorically fed cxC26 mice and ncxC26 mice, with 70 metabolites having significant different contribution (FDR < 0.05) from glucose in at least one tissue between the two groups. The significantly changed metabolites are mainly involved in glycolysis/gluconeogenesis, TCA cycle, amino acid metabolism and nucleotide metabolism. The heatmap shows a generally higher glucose contribution to tissue metabolites in the cxC26 mice than the isocalorically fed ncxC26 mice. This is more vividly shown in the volcano plot comparing the two groups, where almost all the significantly changed contributions lie on the right side of the plot (Figure S6A). In contrast, the heatmap shows a generally lower glucose contribution to tissue metabolites in the isocalorically fed ncxC26 group than the ncxC26 group ad libitum, which is also evident in the corresponding volcano plot (Figure S6B), reflecting normal response of tissue glucose utilization to reduced food intake. Notably, the metabolite with the most significantly increased labeling from glucose was alanine in the cxC26 tumors, suggesting dramatically higher synthesis of alanine in cachectic tumors than non-cachectic tumors (Figure S6C). These results indicate sustained glucose utilization by tissues in the cachectic mice despite reduced food intake while glucose utilization in non-cachectic mice was reduced in response to reduced food intake. The PCA analysis of labeling profiles further revealed the tissues with sustained glucose utilization in the cxC26 mice, and they were gWAT, BAT, kidney, spleen, and small intestine, liver as well as tumor (Figure S6D).

We next focused on glucose contribution to the tissue TCA cycle to examine specifically glucose oxidation by tissues (Figure 3E). The result showed higher glucose contribution to the TCA cycle in BAT, brain, kidney, lung, spleen, small intestine, and tumor in the cxC26 mice compared to the isocalorically fed ncxC26 mice. Comparison with the ncxC26 mice ad libitum revealed that glucose contribution to these tissues dropped in the isocalorically fed ncxC26 mice but showed smaller or even no changes in the cxC26 mice (Figure 3E). Thus, glucose oxidation was sustained in specific tissues in cachectic mice despite reduced food intake. This conclusion is in line with data from the KL and KP models where glucose contribution to tissue TCA was mostly maintained in the cachectic mice compared to their respective ad libitum fed control groups (Figure S7).

### Glucose uptake flux by tumors is an insignificant fraction of total glucose consumption flux in host

One explanation for the negative energy balance that drives weight loss in cachexia could be the high nutrient demand by the tumor. One often-cited nutrient is glucose, as a hallmark of tumor metabolism is the Warburg effect, referring to the high glucose consumption flux by tumors. To determine the extent to which tumor glucose consumption can impact host glucose turnover flux, we measured glucose uptake flux by tumors. Specifically, we administered ^14^C-labeled 2-deoxyglucose (^14^C-2DG) to cxC26 and ncxC26 mice at day 12. We found that the glucose uptake flux of either cxC26 or ncxC26 tumors represented an insignificant fraction of total glucose consumption flux in the whole animal, with the glucose uptake flux of cxC26 tumors (62.1 nmol/min) amounting to only 3.4% of the glucose turnover flux (1817.4 nmol/min) (Figure S8). Although the glucose uptake flux by cxC26 tumors was higher than ncxC26 tumors, the difference (19.3 nmol/min) accounted for only 1.1% of the glucose uptake flux in cxC26 mice. These results show that the tumor’s glucose demand is far from accounting for the changes to the glucose turnover flux at the whole-body level. Of note, tissues such as BAT, lung and spleen had higher glucose uptake flux in cxC26 mice compared to isocalorically fed ncxC26 mice despite similar levels of circulating glucose (Figure S8).

### Increased lactate oxidation by specific tissues in cachectic mice

In addition to glucose, lactate is another major carbohydrate energy nutrient. We aimed to determine how its fluxes change in the C26 model. To quantify lactate turnover flux, we used double-catheterized mice to collect arterial blood from carotid artery catheter while infusing uniformly ^13^C-labeled lactate via jugular vein catheter. This is to avoid the acute stress arising from tail vein blood collection, which can perturb lactate production (Lee et al., 2023). Our results show no change in lactate turnover flux but increased lactate oxidation by specific tissues in cachectic mice compared to the isocaloric non-cachectic mice (Figure 4A). For example, cachectic tumors exhibited the most significant elevation in lactate’s contribution to its TCA cycle (FDR < 0.05). Several tissues showed more than two-fold changes in their TCA contribution by lactate, including soleus (fold-change = 2.5, p = 0.008), heart (fold-change = 2.9, p = 0.013), and BAT (fold-change = 2.7, p = 0.008) (Figure 4B). However, these changes did not remain after the more stringent FDR correction. The labeling of other tissue metabolites was mostly unchanged across tissues (5 among the 559 detected tissue metabolites) (Figure S9A and B). Overall, whole-body lactate turnover flux remained unchanged, but tumor, soleus, heart, and BAT showed increased lactate oxidation in cachectic mice.

**Figure 4:**
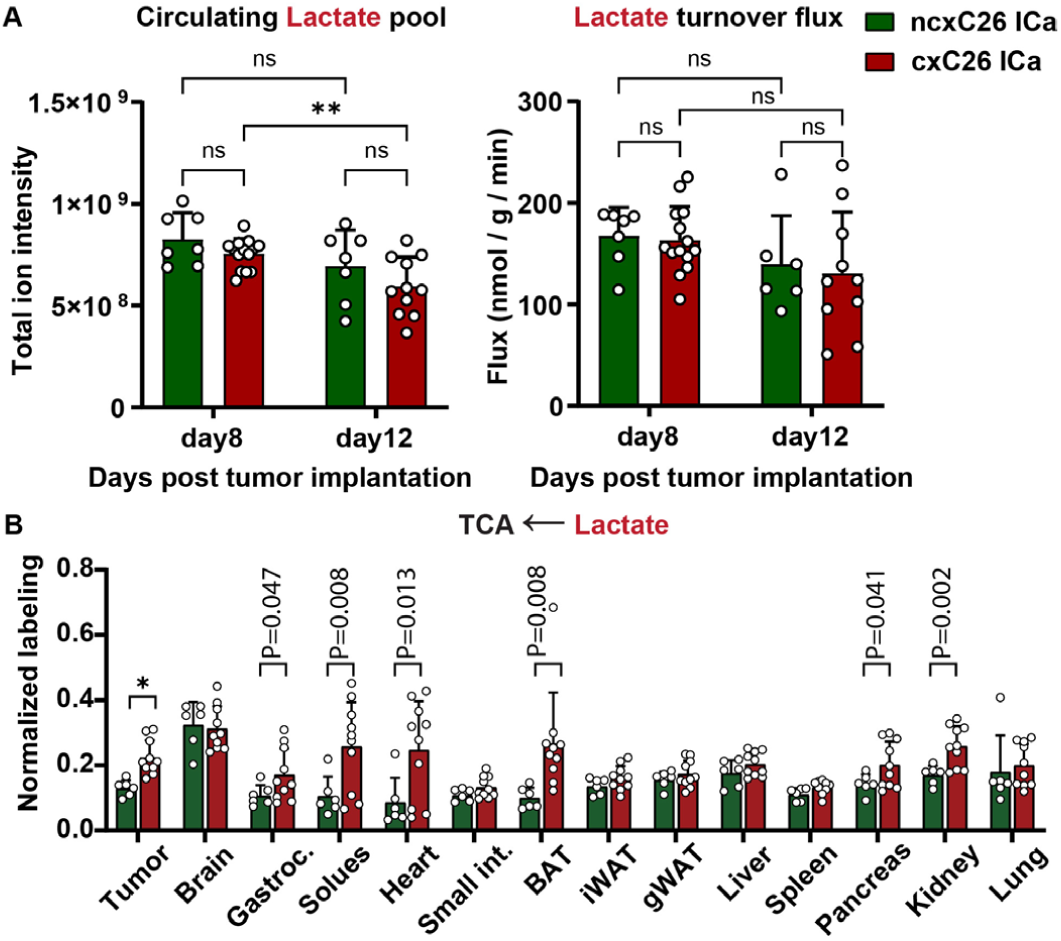
Lactate turnover flux was not altered in cachectic mice but lactate oxidation was increased in specific tissues. (A) Circulating lactate pool and turnover flux in cxC26 and ncxC26 mice under isocaloric feeding at day 8 and day 12. Significance of the differences: ns: non-significance, *P < 0.05, ** P < 0.01, *** P < 0.001 between groups by two-way ANOVA. (B) Normalized labeling of malate in tissues and tumors under ^13^C-lactate infusion at day 12. Significance of the differences: no symbol: not significant, *FDR < 0.05, ** FDR< 0.01, *** FDR < 0.001 between groups by two-tailed t-test with multiple corrections. “P” represents raw p-value (P < 0.05) without multiple correction between groups. Data are shown as mean ± s.d. (n= 6-14)

### Elevated glutamine turnover flux and widespread alterations in glutamine utilization by tissues in cachectic mice

Given the changed glucose metabolism in cachectic mice, next we expanded our nutrient list to include glutamine and alanine, which together with glycerol and lactate are the major gluconeogenesis substrates. We performed uniformly ^13^C-labeled glutamine and alanine infusions. The glutamine pool in cxC26 mice was slightly lower than ncxC26 mice at day 12. However, glutamine turnover flux in cxC26 mice was significantly elevated (by 35%) compared to isocalorically fed ncxC26 mice at day 12, despite both groups consuming the same amount of food (Figure 5A). Additionally, this flux in cxC26 mice was even higher than that in the pre-cachectic stage at day 8. Similarly, alanine turnover flux was also significantly elevated (by 27%) in cxC26 mice compared to ncxC26 mice at day 12 (Figure 5B). These data imply that the maintenance of glucose production flux in cachectic mice was underpinned by augmented gluconeogenic input from glutamine and alanine. To test this hypothesis, we will calculate production fluxes from each of the gluconeogenesis substrates to glucose in a later section.

**Figure 5.**
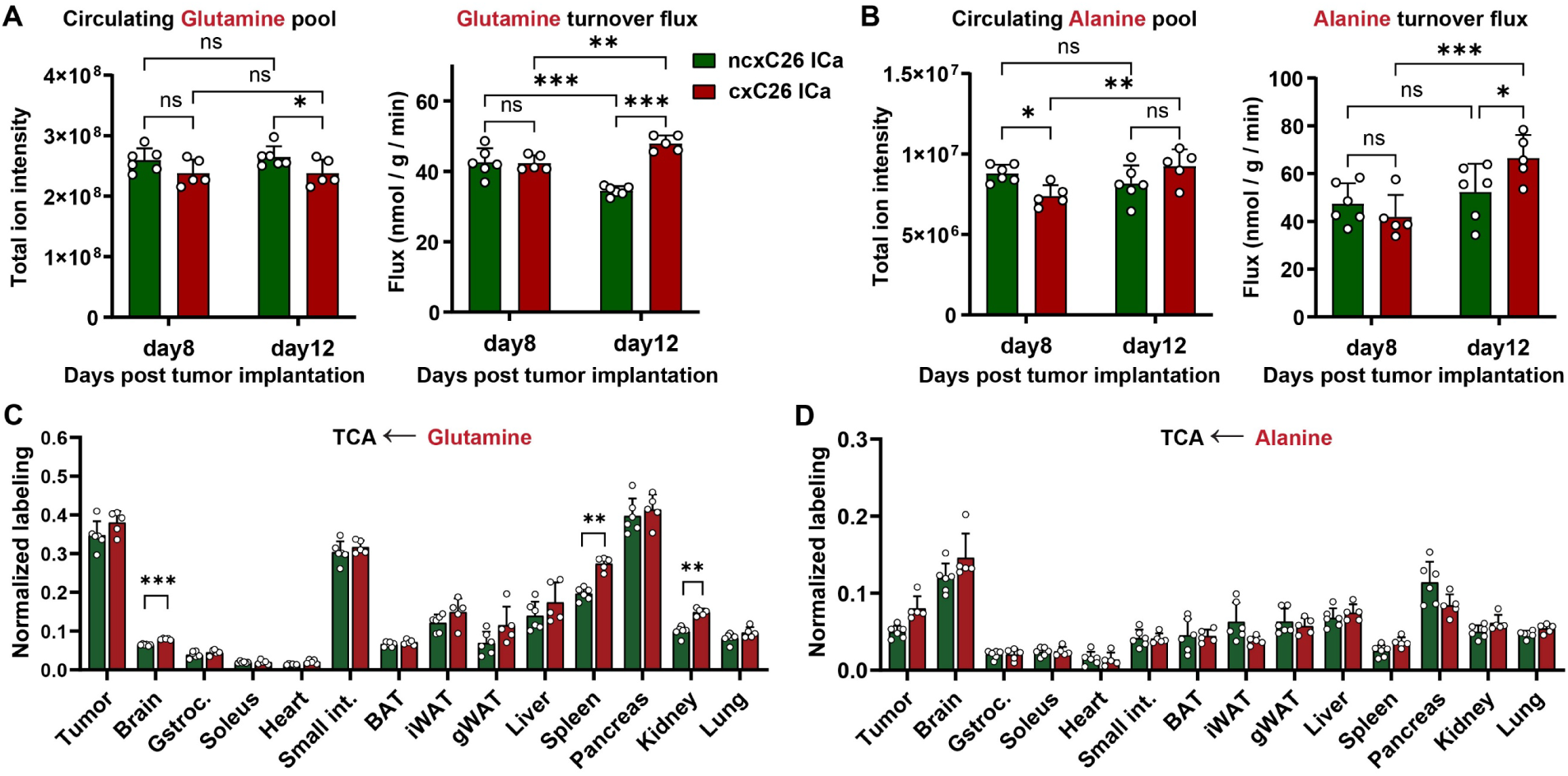
Glutamine and alanine turnover fluxes were elevated in cachectic mice. (A) Circulating glutamine pool and turnover flux in cxC26 and ncxC26 mice at day 8 and day 12 post tumor implantation. (B) Circulating alanine pool and turnover flux in cxC26 and ncxC26 mice at day 8 and day 12. (C and D) Normalized labeling of malate under (C) ^13^C-glutamine infusion and (D) ^13^C- alanine infusion in cxC26 and ncxC26 mice at day 12. Data are shown as mean ± s.d. Significance of the differences: (A, B) ns: non-significance, *P < 0.05, ** P < 0.01, *** P < 0.001 between groups by two-way ANOVA or (C, D) no symbol: not significant, * FDR < 0.05, ** FDR < 0.01, *** FDR < 0.001 between groups by two-tailed t-test with multiple corrections. All data was obtained for mice under isocaloric feeding (ICa). (n= 5-6)

As for the nutrients described above, we determined the contribution of glutamine and alanine to the TCA cycle in tissues. The glutamine contribution was significantly increased in brain, spleen and kidney of cxC26 mice, but there were no significant changes in alanine contribution to tissues compared to food intake-controlled ncxC26 mice (Figure 5C and D). The glutamine contribution to other metabolites was altered in tumor and most tissues except gWAT and small intestine (Figure S10A). A total of 90 tissue metabolites had significantly changed contribution from glutamine in cachectic mice compared to food intake-controlled ncxC26 mice, with 74 having increased contribution and 16 having decreased contribution (Figure S10B). In contrast, the contribution of alanine to tissue metabolites was not significantly changed. Only 4 metabolites showed significant difference between cxC26 mice and food intake-controlled ncxC26 mice (Figure S10C and D). Altogether, the results demonstrate widespread alterations in glutamine utilization in cachectic mice.

### Increased glucose production flux from glutamine and alanine in cachectic mice

In the above, we identified glucose together with its two precursors glutamine and alanine as nutrients with increased turnover fluxes in cachexia, suggesting increased glucose production fluxes from glutamine and alanine. To test this, we calculated the production fluxes from each of the traced nutrients to glucose and to other circulating nutrients using the labeling data and tracer infusion rates (Hui et al., 2020). As illustrated in Figure 6A, a production flux from a nutrient A to a nutrient C represents the direct contribution from A to C, while the abovementioned contribution (as measured by normalized labeling, labeling of C divided by labeling of A) reflects total contribution from A to C including both the direct contribution from A to C and also indirect contribution via another nutrient B. This method involves solving a set of linear mass balance equations (see Methods). The results for glucose are shown in Figure 6B. Production fluxes to glucose from glutamine and alanine were significantly elevated in cxC26 mice compared to food intake-controlled ncxC26 mice. Biochemically glucose cannot be made from fatty acids and the small calculated production flux from palmitate to glucose reflects scrambling of carbon atoms in the TCA cycle (Tetrick and Odle, 2020).

**Figure 6.**
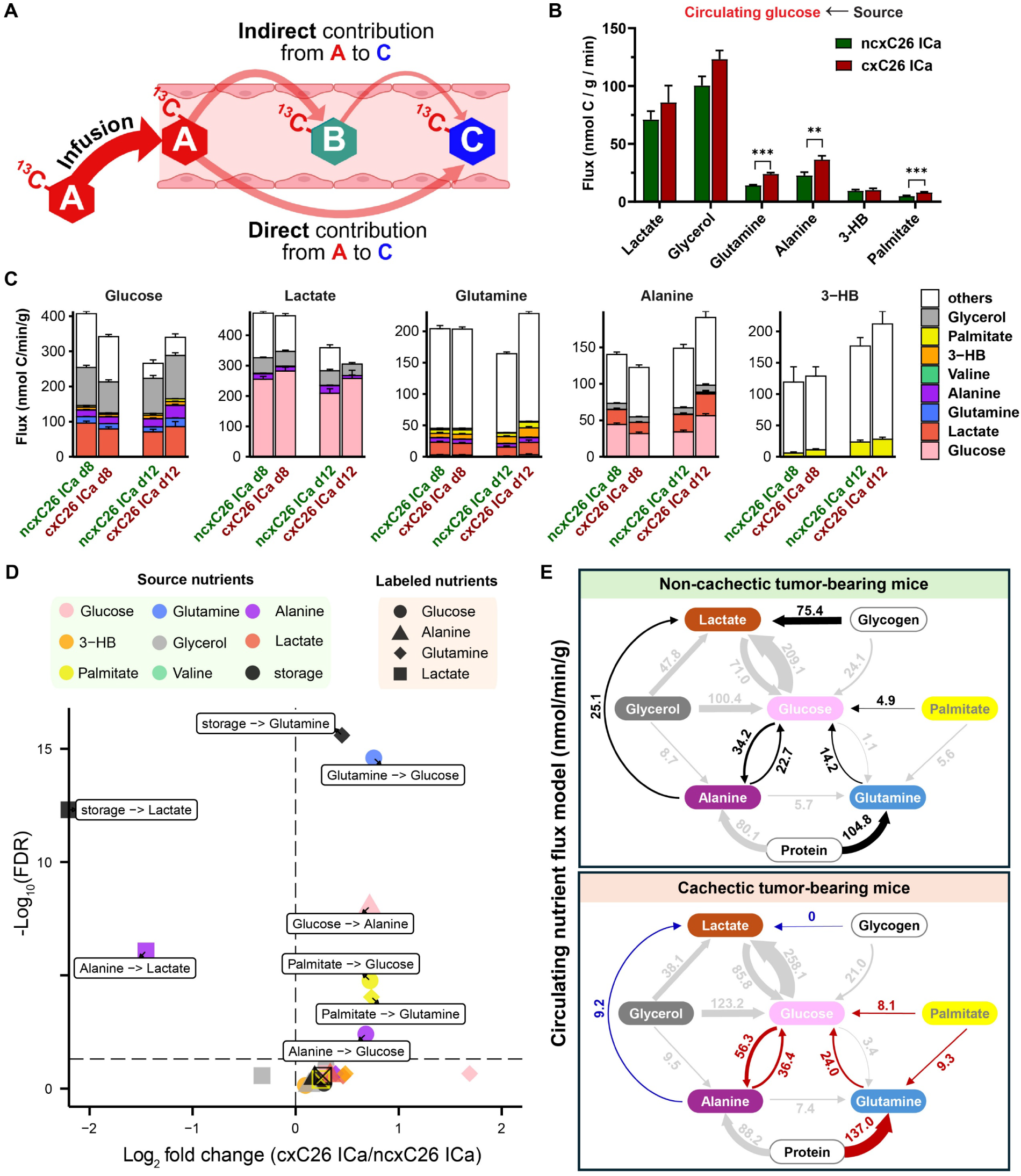
Increased contribution from glutamine and alanine to glucose production and elevated glucose-alanine cycling flux in cachectic mice. (A) Illustration of direct contribution flux (or production flux) to a circulating nutrient from other circulating nutrients. (B) Production flux to circulating glucose from each of other gluconeogenesis substrates in cxC26 and ncxC26 mice at day 12 post tumor implantation. (C) Stacked plots of production flux to each of the circulating nutrients from other circulating nutrients in cxC26 and ncxC26 mice at day 8 and day 12. (D) Volcano plot showing significantly changed interconversion fluxes between circulating nutrients in cxC26 mice compared to ncxC26 at day 12. (E) Interconversion fluxes between circulating nutrients and nutrient storages in cxC26 and ncxC26 mice at day 12. Significantly increased and decreased fluxes are highlighted with red and blue arrows, respectively. Data are shown as mean ± s.e.m. Significance of the differences: no symbol: not significant, *FDR < 0.05, ** FDR < 0.01, ** FDR < 0.001 between groups by two-tailed t-test with multiple corrections. All data was obtained for mice under isocaloric feeding (ICa). (n= 5-11).

In addition to the result for glucose, the production fluxes for lactate, glutamine, alanine, and 3-HB were also calculated, shown in Figure 6C as stacked bars and also in Figure 6D as a volcano plot. The remaining three nutrients, namely valine, palmitate, and glycerol, were either minimally or not produced from other circulating nutrients. Of note, a nutrient can be produced from sources other than the eight nutrients in our list. These other sources can be another circulating nutrient or a nutrient storage (e.g., glycogen), represented as “others” in Figure 6C. Interestingly, the results revealed lactate as another nutrient with significantly changed production. Specifically, in cachectic mice, the production flux from alanine to lactate was significantly lowered, and there was no lactate production from sources other than the nutrients we probed in this study. This was not the case in ncxC26 mice or pre-cachectic mice, where a substantial production flux originated from “other” sources which are presumably muscle glycogen. Therefore, this result suggests impaired lactate production from muscle glycogen in cachectic mice.

### Elevated glucose-alanine cycling flux and glutamine production flux from tissue proteins

Combining the production fluxes to each of the nutrients, we can construct a flux model describing interconverting fluxes between these nutrients. The glucose-related parts of the flux models for cachectic and non-cachectic mice are shown in Figure 6E. To determine the flux from storage to glucose, we estimated fluxes from unmeasured free fatty acids (FFAs) and subtracted them from the flux from “others” (Hui et al., 2020) (Methods). The resulting storage flux can be taken as the conversion flux of glycogen to glucose. Similarly, we obtained the storage flux from protein to glutamine and alanine, respectively. As expected from our results above, the storage flux to lactate was suppressed. The flux models additionally uncovered a pronounced upregulation of glucose-alanine cycling, signifying enhanced inter-organ nitrogen shuttling in the cachectic mice. Moreover, the flux models showed higher flux from protein to glutamine, indicating tissue proteins as the source of elevated glutamine turnover flux in cachectic mice.

### Altered fuel preference by specific tissues in cachectic mice

In the above, we have analyzed the contribution from each of the traced nutrients to tissue TCA cycle. The contribution was calculated as the ratio of the TCA labeling to the labeling of the traced nutrient in circulation. As illustrated in Figure 7A, this quantity reflects the total contribution from the circulating nutrient to the tissue TCA cycle, comprising both direct contribution and indirect contribution via other nutrients. To determine fuel preference by tissues, we calculated the direct contribution from the circulating nutrients to the TCA intermediates malate and succinate in tissues (Figure 7B and 7C) (see Methods). The results revealed a largely stable fuel preference across tissues and nutrients in cachectic mice, with a small number of changes. The most significantly changed tissue-nutrient relations were spleen and kidney’s usage of glutamine and cachectic tumors’ usage of glucose. Glucose was also used more by spleen but less by kidney. In addition, lactate was used more by soleus, BAT, and heart, and alanine was used less by soleus, gastrocnemius, BAT, iWAT and pancreas. Thus, our results revealed changes in tissue fuel usage in cachectic mice.

**Figure 7.**
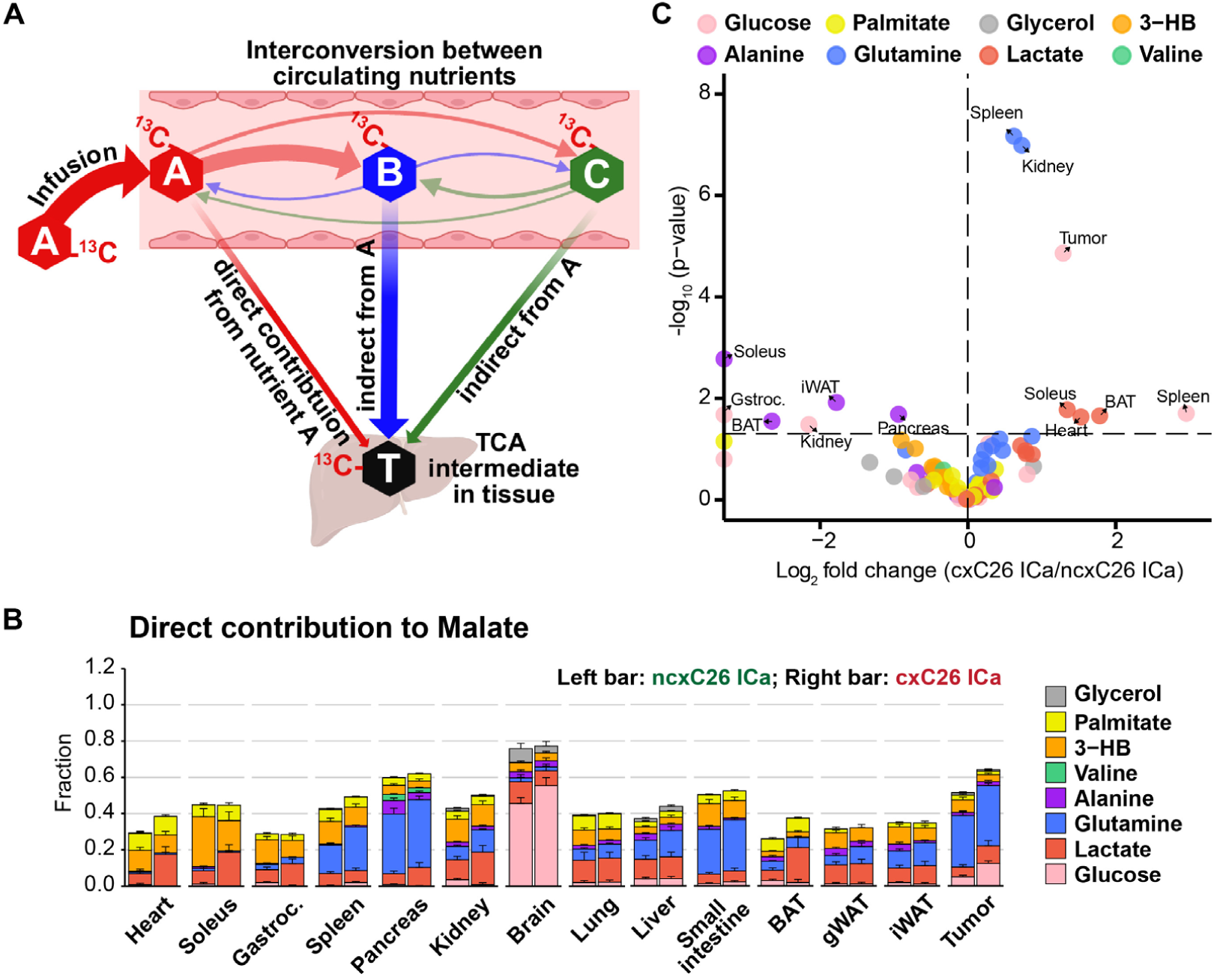
Fuel preference was altered in specific tissues in cachectic mice. (A) Illustration of direction contribution of a circulating nutrient to tissue TCA cycle. (B) Stacked bar plot of fuel preference (direction contribution of circulating nutrients to malate) of tissues and tumors in cxC26 and ncxC26 mice under isocaloric feeding (ICa), at day 12 post tumor implantation. (C) Volcano plot of the same data as in (A). All data are shown as the mean ± sem. (n= 5-11)

## Discussion

Our study presents a comprehensive flux quantification of all major circulating nutrients in a widely used mouse model of cancer cachexia under highly controlled conditions. The flux results provide direct and comprehensive answers to the fundamental question of how host metabolism is altered in cancer cachexia. We find that energy metabolism is largely preserved in lipid and protein metabolism and altered in glucose-related metabolism. Importantly, the metabolic alterations we identified are not simply due to changed energy balance but reflect fundamental dysregulations of those pathways caused more directly by tumors.

Cancer cachexia is viewed as a complex metabolic disorder, often described with a long list of metabolic perturbations (Berriel Diaz et al., 2024). Our study presents a more simplistic picture, with stable fluxes in lipid and protein metabolism. This contradiction is in part due to our highly controlled experimental system. First, we had mice bearing non-cachectic tumors to control for metabolic changes arising from tumor mass as opposed to the cachectic process. Second, we controlled for food intake, which is a potent and broad influencer of metabolism. Third, we were able to eliminate other potential confounders of metabolism (i.e., body weight and composition, energy expenditure, and energy excretion) as cachectic and non-cachectic mice had similar values for these variables under our isocaloric feeding protocol. Thus, the changes revealed by our flux results are cachexia-inherent, which are constrained to metabolism of glucose and related nutrients.

A major finding in our study is the impaired response of glucose metabolism to reduced food intake in cancer cachexia. In mice without cancer cachexia, food restriction lowers blood glucose levels and suppresses glucose turnover flux (i.e., rates of glucose production and consumption). This response is lost in cachectic mice, where glucose turnover flux remains high despite reductions in food intake. For example, food restriction lowers glucose oxidation in the tumor and BAT of non-cachectic mice, but these remain high in the cachectic mice (Figure 3E). Moreover, glucose oxidation in the small intestine, spleen, pancreas, and lung are significantly higher in the cachectic mice as compared to the isocaloric non-cachectic mice. Glucose oxidation can be controlled by both insulin-dependent and independent mechanisms. Since insulin levels were similar between cachectic and isocaloric non-cachectic mice (Figure S11), we speculate that insulin-independent pathways, such as catecholamine signaling, or expansion of cell populations with constitutive (GLUT1- or GLUT3-mediated) glucose uptake may underlie the elevated glucose utilization observed. Immune cells are an especially intriguing candidate, given their high glycolytic rates and expansion during states of systemic inflammation, including C26-induced cachexia (Petruzzelli et al., 2022).

The continuous consumption of glucose by cachectic tumors, even in the food restricted state, may suggest that these tumors predispose to cachexia by increasing energy expenditure (i.e., “hypermetabolism”) or depriving other tissues of glucose. Indeed, cachectic tumors have higher [^18^F]-FDG uptake than non-cachectic tumors in human patients (Olaechea et al., 2022) and preclinical models (Burvenich et al., 2024; Penet et al., 2011). However, our data does not support this “hypermetabolism” theory. First, there is no difference in total energy expenditure when comparing isocalorically matched cachectic and non-cachectic mice (Liang et al., 2025; Queiroz et al., 2022; Verlande et al., 2022). Second, the relative consumption of glucose by the tumor in comparison to whole body glucose turnover flux is quite low. The difference between isocalorically matched cachectic and non-cachectic mice amounts to ∼1% of total glucose turnover flux. Therefore, we conclude that glucose consumption by cachectic tumors is unlikely a major contributor to energy deficit during cachexia.

The high glucose consumption flux during cachexia is balanced by a proportionally high glucose production flux. Our study identifies alanine and glutamine as substrates that are responsible for the sustained glucose production (Figure 6B). Alanine is a well-known gluconeogenic precursor and we observed an increase in glucose-alanine cycling flux. Specifically, cachectic tumors have high glucose to alanine flux (Figure S6C), which may serve to dispose of excess nitrogen while recycling carbon skeletons to the liver for glucose production. While alanine contributes meaningfully to glucose carbon recycling, glutamine is quantitatively more important for transporting protein-derived carbon to the glucose pool (Nurjhan et al., 1995). We observed a significant increase in the flux of stored nutrients to glutamine in cachectic mice (Figure 6D). We also noted increased glutamine oxidation by the spleen and kidney (Figure 5C) and glutamine convertion to glucose by a currently undefined tissue (Figure 6B). Given its known preference for glutamine as a gluconeogenic substrate, the kidney may play a central role in this process (Stumvoll et al., 1998). Future experiments will attempt to restore normal glucose homeostasis (e.g., suppressing elevated gluconeogenesis) in cachectic animals and determine whether there are ameliorating effects on cachexia (Liu et al., 2025).

The sustained glucose turnover flux and elevated metabolic cycling must impart a caloric cost on the host, and it is unclear why this cost does not appear as measurable energy expenditure. One potential explanation is that there exists a metabolic tradeoff where one form of energy expenditure is replaced or offset by another. Cachectic mice and humans are known to have low physical activity (Bruggeman et al., 2016; Counts et al., 2020), and the normal increase in physical activity that occurs with food restriction is missing in cachectic animals including our C26 mice (Liang et al., 2025; Queiroz et al., 2022). Therefore, cachectic mice “save” energy compared to the food intake-matched control group, and this saved energy can be “re-routed” to support other processes (e.g., immune system activation) that are necessary for surviving cancer. Future work can test this potential link.

### Limitation of the study

Several limitations of the present study merit consideration. First, our analyses were confined to the post-absorptive (fasted) state, precluding examination of nutrient flux dynamics under fed conditions, wherein distinct metabolic phenotypes may emerge. It remains possible that dietary intake could differentially modulate substrate utilization in cachectic versus non-cachectic states, particularly in regard to postprandial lipid and protein metabolism. Second, although our isotopic tracer panel encompassed eight major circulating nutrients relevant to energy metabolism, it did not interrogate other critical metabolic domains such as one-carbon metabolism, redox homeostasis, and nucleotide biosynthesis, which may be perturbed in cancer cachexia. Third, the variability observed in lactate flux measurements—despite use of double-catheterized animals and arterial sampling—is likely attributable to lactate’s high sensitivity to acute adrenergic stress (Lee et al., 2023), underscoring the technical challenges inherent in capturing its physiological flux with precision.

Furthermore, while our findings were corroborated in two independent genetically engineered models of lung cancer cachexia, the bulk of our quantitative flux analyses were conducted in the C26 colon carcinoma model. Thus, although the observed glucose-centric metabolic remodeling appears robust, its universality across diverse tumor types and cachexia subtypes warrants further empirical validation. Additionally, our study exclusively utilized male mice. Given established sex differences in metabolic regulation and tumor biology, it is imperative that key findings be validated in female cohorts to ensure biological generalizability. Lastly, while our analysis revealed profound shifts in systemic carbohydrate metabolism, the upstream regulatory mechanisms—whether tumor-secreted factors, host immune responses, or endocrine perturbations—remain to be elucidated. Future work aimed at mechanistically decoding these regulatory circuits will be essential for translating these metabolic insights into therapeutic strategies.

## STAR★METHODS

## KEY RESOURCES TABLE

| REAGENT or RESOURCE | SOURCE | IDENTIFIER |
| --- | --- | --- |
| **Chemicals, peptides, and recombinant proteins** |  |  |
| 2-Deoxy-D-glucose ($^{14}\text{C}_6$) | PerkinElmer | NEC720A050UC |
| D-glucose ($^{13}\text{C}_6$, 99%) | Cambridge Isotope Laboratories | CLM-1396 |
| Sodium L-lactate ($^{13}\text{C}_3$, 98%)<br>20% w/w In $\text{H}_2\text{O}$ | Cambridge Isotope Laboratories | CLM-1579 |
| L-alanine ($^{13}\text{C}_3$, 99%) | Cambridge Isotope Laboratories | CLM-2184-H |
| L-glutamine ($^{13}\text{C}_5$, 99%) | Cambridge Isotope Laboratories | CLM-1822-H |
| glycerol ($^{13}\text{C}_3$, 99%) | Cambridge Isotope Laboratories | CLM-1510 |
| Sodium D-3-hydroxybutyrate<br>($^{13}\text{C}_4$, 99%) | Cambridge Isotope Laboratories | CLM-3853 |
| Potassium palmitate<br>($^{13}\text{C}_{16}$, 98%) | Cambridge Isotope Laboratories | CLM-3943 |
| L-valine ($^{13}\text{C}_5$, 99%) | Cambridge Isotope Laboratories | CLM-2249-H |
| DMEM with 4.5 g/L glucose | Corning | 10-013-CV |
| Penicillin & Streptomycin | HyClone | SV30010 |
| Fetal Bovine Serum | R&D SYSTEMS | S11550H |
| Normal Saline | HyClone | Z1376 |
| EcoLume™ Liquid Scintillation<br>Cocktail | MP Biomedicals | 0188247001 |
| Antifoam B Emulsion | Sigma-Aldrich | A5757 |
| Bovine serum albumin, fatty<br>acid free, 96% | Sigma-Aldrich | A6003 |
| glycerokinase | Sigma-Aldrich | G6278 |
| **Critical commercial assays** |  |  |
| Mouse/Rat Metabolic Hormone<br>Discovery Assay® Array | Eve Technologies | RDMET12 |
| **Experimental models: Cell lines** |  |  |
| cxC26 cells | (Bonetto et al., 2016) |  |
| ncxC26 cells | (Brattain et al., 1980) |  |
| **Experimental models: Organisms/strains** |  |  |
| Mouse: CD2F1, male | Charles River laboratory | 033CDF-1 |
| Mouse: KP ($\text{Kras}^{\text{LSL-G12D/+}}$;<br>$\text{Lkb1}^{\text{flox/flox}}$) | (Bhatt et al., 2019) | |
| Mouse: KL ($\text{Kras}^{\text{LSL-G12D/+}}$;<br>$\text{p53}^{\text{flox/flox}}$) | (Guo et al., 2013) | |
| **Recombinant DNA** |  |  |
| Lentiviral-Cre | UI Viral Vector Core | VVC-U of Iowa-5 |
| **Software and algorithms** |  |  |
| EL-MAVEN software | Elucidata | https://docs.polly.elucidata.io/index.html |
| Rstudio | Posit | https://posit.co/download/rstudio-desktop/ |
| AccuCor | Github | https://github.com/XiaoyangSu/AccuCor |
| Prism 10 | Graphpad | https://www.graphpad.com/ |
| Excel | Microsoft | https://www.microsoft.com/en-us/microsoft-365/excel |
| Biorender | Biorender | https://www.biorender.com/ |
| **Other** |  |  |
| Jugular Vein Catheters for Mice | Instech Laboratories | C20PUMJV1301 |
| Carotid Artery Catheters for Mice | Instech Laboratories | VAMB1B |
| One Channel Vascular Access Button™ for Mice | Instech Laboratories | VABM1B/25 |
| Two Channel Vascular Access Button™ for Mice | Instech Laboratories | VABM2VB/25R25 |
| chow diet | PicoLab Mouse Diet 20 | 5053 |
| buprenorphine base ER 0.5 mg/mL | Wedgewood | Rx # 7270862 |
| Microvette® CB 300 Serum CAT | Sarstedt | 16.440.100 |
| Microvette® 500 EDTA K3E | Sarstedt | 20.1341.102 |
| XBridge BEH Amide XP column | Waters | 176002889 |

## EXPERIMENTAL MODEL

### Cell lines

The cachectic C26 (cxC26) and non-cachectic C26 (ncxC26) cells were a kind gift from Andrea Bonetto (Bonetto et al., 2016) and Nicole Beauchemin, respectively. Nicole Beauchemin obtained the ncxC26 from Brattain MG, who established the C26 (also named CT26 or colon carcinoma 26) cell line (Brattain et al., 1980). All C26 cells were maintained using high glucose DMEM (Corning) with 10% fetal bovine serum (FBS, R&D Systems) and 1% penicillin/streptomycin (Hyclone) at 37°C under 5% CO^2^ in a humidified atmosphere.

### Animals

Animal care and experimental procedures were conducted with the approval of the Institutional Animal Care and Use Committees (IACUC) of Harvard Medical School and Harvard T.H. Chan School of Public Health. CD2F1 (BALB/c x DBA/2) male mice were purchased from Charles River and used for this study when the mice were 13 – 18 weeks old. All mice were housed individually using white colored paper bedding (ALPHA pad®, Shepherd Specialty Papers) with paper nesting materials (Enviro-dri®, Shepherd Specialty Papers) to facilitate accurate measurement of food leftover on the bedding. Daily food intake was assessed by weighing the change in food pellets on the cage lid, subtracting any food left on the bedding. Mice cages were maintained at room temperature (22°C) with a 12-hour light-dark cycle, and mice were fed either ad libitum or controlled isocaloric feeding with standard chow diet (PicoLab 5053, LabDiet).

For tumor implantation, cxC26 and ncxC26 cells were collected, washed and resuspended in saline, and one million cells in 200 µl of saline were injected into the right flank of mice under anesthesia. Body weight was recorded bi-daily at 9 AM following tumor implantation. For isocaloric feeding, the precise daily food amount from day 8 to 12 post-tumor implantation was based on the food intake of cxC26 mice ad libitum, specifically: day 8: 3.1g, day 9: 2.8g, day 10: 1.7g, and day 11: 1.1g, as illustrated in Figure S1. This daily food allocation was divided into 3 equal portions and dispensed every 4 hours during the night at 7 PM, 11 PM and 3 AM) using an automatic feeder (Figure 1B). Leftover food on the bedding was monitored daily and returned to the cage if necessary. Mice that did not consume all their food (more than 1 gram of cumulative leftover during the entire period of isocaloric feeding) were excluded from experiments.

Animal experiments using *Kras^LSL_G12D/+^; p53^flox/flox^* (KP) and *Kras^LSL-G12D/+^; Lkb1^flox/flox^* (KL) non-small lung cancer cachexia models were performed following protocols approved by the Rutgers IACUC. Lung tumor induction was performed as previously described (Bhatt et al., 2019; Guo et al., 2013). Briefly, tumors were induced in 8-10 weeks old mice via intranasal administration of Lentiviral-Cre (University of Iowa Viral Vector Core) at 4 x 10^7^ plaque-forming units (pfu) per mouse in KP mice, and 2.5 x 10^7^ pfu per mouse in KL mice. Food intake of KP mice was obtained using metabolic cages (Oxymax-CLAMS) post 13 weeks infection.

## METHOD DETAILS

### Isotope tracer infusion

For stable isotope tracing, the jugular vein catheter (C20PUMJV1301, Instech Laboratories) was implanted into the right jugular vein of mice. To minimize potential impact to systemic metabolism by the catheterization procedure, jugular vein catheters were implanted in mice two weeks before cancer cell injection. The catheter was connected to a single-channel button (VABM1B/25, Instech Laboratories) placed subcutaneously on the back of mice. For lactate infusion, an additional carotid catheter (C10PU-MCA2A09, Instech Laboratories) was implanted into the left carotid artery and connected to a two-channel button (VABM2VB/25R25, Instech Laboratories) under the skin alongside the jugular vein catheter. Catheters were filled with heparinized saline (20 U/ml) as a lock solution. Surgery procedures were conducted under aseptic conditions, with anesthesia using isoflurane to ensure animal welfare and minimize pain. Post-surgery, each mouse received a subcutaneous injection of 50 µl of analgesic buprenorphine (0.5 mg/mL) into the left flank to effectively manage postoperative pain. Mice that have undergone surgery were allowed a recovery period of 1-2 weeks before being used in any additional experiments.

Mice were fasted for 5 hours before tracer infusion. The catheter was connected via a swivel-tether system to a tracer-filled syringe that was pushed by a syringe pump, as described previously (Hui et al., 2017). Each ^13^C-labeled tracer was infused for 2.5 hours at a rate of 0.1 µL/g/min on day 8 and day 12 post tumor implantation, with the exception of palmitate, which was infused for 1.5 hours at a rate of 0.2 µL/g/min. The concentration of each tracer was as follows: 200 mM ^13^C-glucose, 378.14 mM ^13^C- lactate, 75 mM ^13^C-glutamine, 30 mM ^13^C-alanine, 15 mM ^13^C-valine, 9.5 mM ^13^C-potassium palmitate (with BSA), and 109.4 mM ^13^C-glycerol. Additionally, the ^13^C-3- Hydroxybutyrate tracer was infused at a concentration of 50 mM on day 8 and 120 mM on day 12. All ^13^C-labeled tracers used were uniformly labeled as all carbon atoms. Blood samples were collected using serum collection tubes (Microvette® CB300, Sarstedt) at the end of infusion from tail without stopping the infusion. For mice infused with lactate, arterial blood was collected via the artery catheter without disturbing the mice, followed by tail blood collection. Blood samples were centrifuged for 10 minutes at 2,000g at 4°C to obtain serum. Mice were euthanized by cervical dislocations shortly after blood collection, and tissues were immediately dissected, quickly wrapped with aluminum foil and clamped using a pre-chilled Wollenberger clamp, and placed in liquid nitrogen. At end of experiments, samples were transferred to -80°C for storage.

### Metabolite extraction from serum and tissue samples

For metabolite extraction of serum samples, 45 µl of pre-cooled extraction buffer (40:40:20, methanol: acetonitrile: water, *v:v:v*, -20°C) was added to 5 µl of serum. For tissue metabolite extraction, frozen tissues were first ground into powder using a CryoMill (Retsch) at cryogenic temperature with liquid nitrogen cooling. ∼1 mL of pre-cooled extraction buffer was then added to 15-25 mg of ground tissue to achieve a final concentration of 25 mg/ml. In the next step, both serum and tissue samples were vortexed for 10 seconds to ensure thorough mixing and then incubated on ice for 10 minutes. Following incubation, the mixture was centrifuged at 1,000g for 10 minutes at 4°C. The supernatant underwent a second centrifugation at 16,000g for 20 min at 4°C. Finally, clear supernatant was transferred to LC-MS vials for subsequent analysis.

### Metabolite measurements by LC-MS

Metabolite extracts were analyzed using a quadrupole-orbitrap mass spectrometer coupled with hydrophilic interaction liquid chromatography (HILIC) as the chromatographic technique. Chromatographic separation was achieved on an XBridge BEH Amide XP Column (2.5 µm, 2.1 mm × 150 mm) with a guard column (2.5 µm, 2.1 mm X 5 mm) (Waters, Milford, MA). Mobile phase A consisted of water: acetonitrile at 95:5 (*v:v*), and mobile phase B consisted of water: acetonitrile at 20:80 (*v:v*), with both phases containing 10 mM ammonium acetate and 10 mM ammonium hydroxide. Separation was conducted under the following linear elution gradient: 0 ∼ 3 min, 100% B; 3.2 ∼ 6.2 min, 90% B; 6.5. ∼ 10.5 min, 80% B; 10.7 ∼ 13.5 min, 70% B; 13.7 ∼ 16 min, 45% B; and 16.5 ∼ 22 min, 100% B, with a flow rate of 0.3 mL/ min. The autosampler and column were maintained at 4°C and 30°C, respectively. The injection volume was 5 µL and the needle was washed between samples using methanol: acetonitrile: water at 40: 40: 20 (*v:v:v*). Mass spectrometry analysis was performed using either a Q Exactive HF or Orbitrap Exploris™ 480 (Thermo Fisher Scientific, San Jose, CA), with mass range from 70 to 1000 *m/z* and polarity switching mode at a resolution of 120,000. Metabolite identification was based on accurate mass and retention time using the EI-Maven (Elucidata) with an in-house library. Natural abundance correction for ^13^C was conducted in R using AccuCor package (Su et al., 2017).

### Measurement of glycerol in serum samples

Glycerol measurement involved derivatization to glycerol-3-phosphate using glycerol kinase. 5 µl of serum sample were mixed with 45 µl of reaction mixture (25 mM Tris, 50 mM NaCl, 10 mM MgCl2, 1 mM ATP, and ∼0.2 U/ml of glycerol kinase in H2O). The mixture was incubated for 15 minutes at room temperature. Following incubation, 200 µl of pre-cooled methanol (-80°C) were added to the mixture, which was then centrifuged at 16,000g for 10 minutes. Supernatant was transferred to a new tube and dried using a Speed Vac at 40°C for 2 hours. The dried residue was dissolved in 50 µl of pre-cooled extraction buffer (40:40:20, methanol: acetonitrile: water, *v:v:v*, -20°C) and transferred to an LC-MS vial for analysis with the HILIC method. Signal intensity of glycerol-3-phosphate was adjusted by subtracting background level of free glycerol-3-phosphate in serum and any contamination from reaction mixture, utilizing procedural control.

### Glucose uptake assay

Mice were fasted for 6 – 8 hours prior to experiment. To minimize urination during experiment, the lower abdomen of each mouse was gently massaged to induce bladder emptying 45 minutes before the 2-DG injection. 1 µCi of ^14^C6-2-Deoxy-D-glucose (PerkinElmer, NEC720A050UC) in saline was injected intraperitoneally (IP). After 45 minutes of injection, the mouse was euthanized and all major tissues, tumor, blood, and urine were collected. Each of the samples (less than 300 mg) was digested using 600 µl of 1.7M potassium hydroxide (KOH) at 70°C for 2 hours. The samples were then allowed to cool at room temperature before 300 µl of isopropanol containing 2M acetic acid and 1% Tween 80 were added to each of them. Subsequently, 400 µl of each digested sample were transferred to a scintillation vial which contained 200 µl of 50% hydrogen peroxide with 15% antifoam B emulsion. The samples were incubated at room temperature for 30 minutes, followed by incubation at 70°C for 20 minutes to fully decompose the hydrogen peroxide and decolorize the samples. Afterward, the samples were allowed to cool at room temperature before 100 µl of 12M acetic acid and 4 mL Ecolume scintillation liquid (MP Biomedicals) were added to each vial. The samples were vigorously vortexed before being loaded to a HIDEX300S

Liquid scintillation counter (Turku, Finland) for radioactive carbon-14 (^14^C) measurement.

### Measurement of insulin in plasma samples

Blood samples were collected using K3EDTA-coated plasma collection tubes (Microvette® 500 EDTA K3E, Sarstedt), with protease inhibitor cocktails and DPPIV added immediately. Plasma was then isolated by centrifugation at 2,000g for 10 min at 4°C. Insulin in the plasma was quantitated using Mouse/Rat Metabolic Hormone Discovery Assay^®^ Array (Eve Technologies).

### Turnover flux of circulating nutrients

The turnover flux of a circulating nutrient was calculated using the following equation

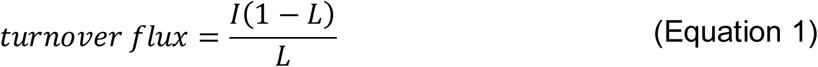

where is 𝐼 the tracer infusion rate and 𝐿 is the labeled fraction of the fully labeled form of the infused uniformly ^13^C-labeled tracer (Hui et al., 2017).

### Calculation of production fluxes of a circulating nutrient from other circulating nutrients

The production fluxes of a circulating nutrient from other circulating nutrients were calculated using equations that we described in a previous study with further details of the calculation available in that publication (Yuan et al., 2025). To explain the calculation here, we use the calculation of glucose production fluxes as an example. Under nonperturbative steady-state ^13^C-glucose infusion with an infusion rate *I*, the total glucose production flux (*P^total^,* including both ^12^C and ^13^C) is equal to summation of all incoming fluxes to glucose from the 7 other circulating nutrients (lactate, alanine, glutamine, glycerol, palmitate, 3-hydroxybutyrate, and valine) and from glucose tissue storages (𝐹_𝑠_) such as glycogen, or

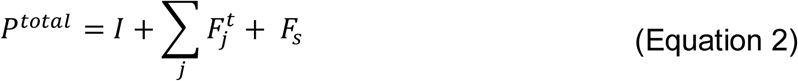

where 𝐹_𝑗𝑗_ is the glucose production flux from circulating nutrient *j*. A second mass balance equation can be written for the labeled glucose pool, which states that the sum of incoming ^13^C fluxes to glucose from different labeled nutrients is equal the outgoing labeled glucose flux, or

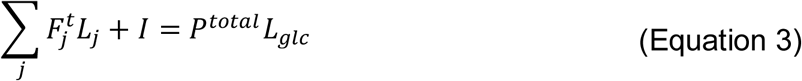

The total glucose production (^12^C and ^13^C) from nutrient *j* can be expressed in terms of the endogenous (^12^C) glucose production from nutrient *j* (𝐹_𝑗_) and its labeling 𝐿_𝑗_, i.e.,

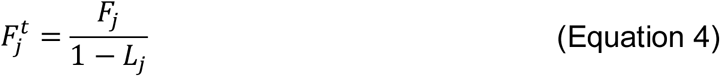

Combining Equations 2 and 3 can generate an equation with P^total^ eliminated. Replacing 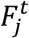 in this equation using Equation 4 yields an equation with the 7 𝐹_𝑗_’s and 𝐹_𝑠_ as the only unkowns. To solve for these unknowns, in addition to one equation for the glucose tracer infusion, one such equation can be written for the infusion of each of the other 7 tracers but with 𝐼 = 0. Integration of the 8 equations from all tracing experiments creates a linear system 𝐴 ⋅ 𝑓 = 𝑏 as shown below

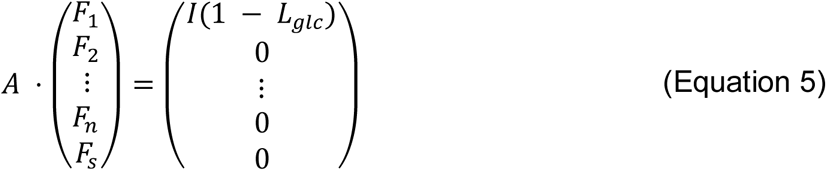

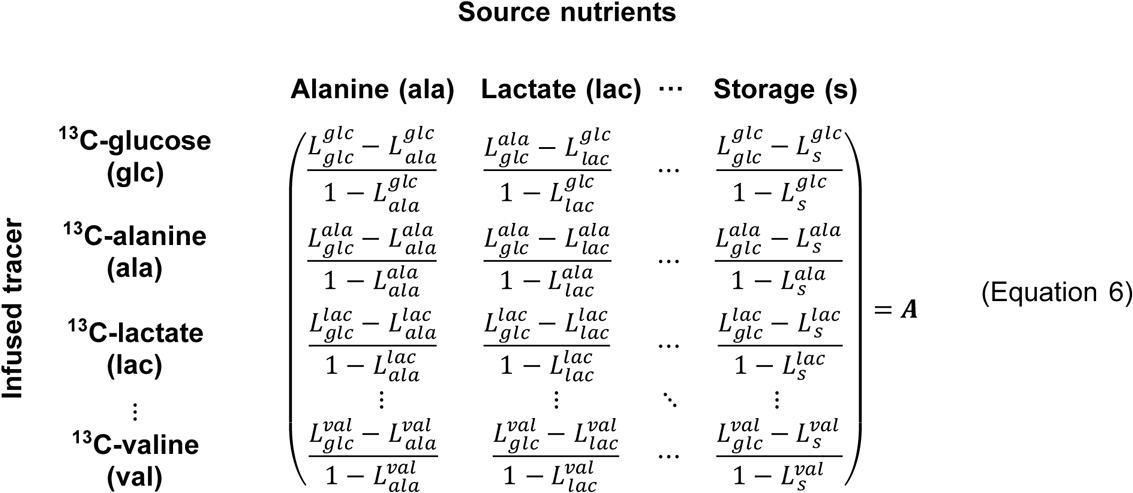

where the storage labeling 𝐿_𝑆_ is assumed negligible, with 𝐿_𝑆_ = 0. In matrix 𝐴, each row corresponds to a different ^13^C-tracing experiment, and each column corresponds to a different production source (circulating nutrients or storage) to glucose. We employed the R package LimSolve to solve the linear algebra, s.t. 𝑓 ≥ 𝟎. To estimate errors, we conducted Monte Carlo simulation by running the matrix optimization 1,000 times. In each simulation, for each measured labeling 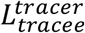, a random value was sampled from a normal distribution with its mean and standard deviation being the mean and standard error of this measured quantity. Each simulation resulted in a computed 𝑓 vector. The standard deviation of 𝑓 was calculated as the standard error of the contribution fluxes from different sources to glucose production. Here we have illustrated our calculation using glucose production as an example. Calculation of the production fluxes to other circulating nutrients were computed in like manner.

### Calculation of flux from storage to a circulating nutrient

To determine the flux from storage to a circulating nutrient such as glucose, alanine or glutamine, we subtract the estimated total FFA flux (to the nutrient) from the remaining flux (labeled as ‘others’ in Figure 6C). The total FFA flux to a nutrient was estimated by dividing the measured palmitate flux to that nutrient by the ratio of palmitate abundance to total FFA abundance in the circulation (0.2111) (Hui et al., 2020).

### Calculation of direct contributions to tissue TCA cycle from circulating nutrients

Direct contributions of circulating nutrients to malate or succinate in tissues were calculated using a slightly modified version of the equation described in our previous study, with detailed calculation procedures available in that earlier publication (Yuan et al., 2025). The contribution fraction of a circulating nutrient 𝑗 to the target nutrient (*TN*) in a tissue is defined as 𝑓_𝑗_. A mass balance equation of ^13^C gives

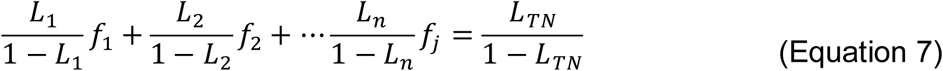

where *L_TN_* is the ^13^C labeling in *TN*. Integrating 8 equations from the 8 ^13^C-tracer infusions gives the following set of equations

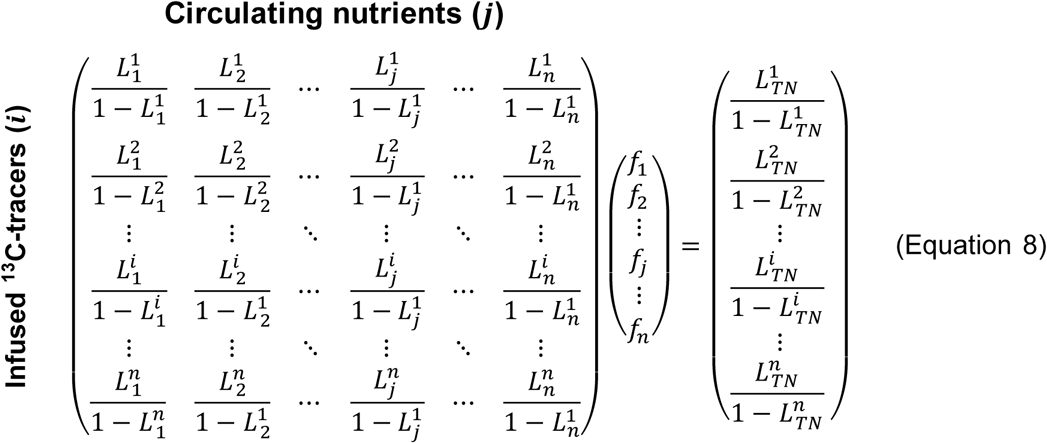

where 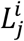 represents the labeling of a circulating nutrient 𝑗 under infusion of ^13^C-tracer 𝑖. The optimization procedure for determining the values of 𝑓’s and their error estimation were performed as described above.

### Statistical analysis

All statistical analyses were performed using GraphPad Prism 10 or R studio. Quantitative data are reported as mean with standard deviation (s.d.). Two-group comparisons were analyzed by a two-tailed Student’s t-test, and more than two-group comparisons were analyzed by One-way ANOVA or Two-way ANOVA. Adjusted p-values (FDR) obtained for multiple comparisons using the Benjamini–Hochberg method. For all analyses, a p-value of < 0.05 was considered significant *(*p < 0.05, **p < 0.01, and ***p < 0.001*). The R package ComplexHeatmap was used to generate the heatmap for the labeling of tissue metabolites by circulating glucose.

## ACKNOWLEDGMENTS

We would like to thank Dr. Andrea Bonetto (cxC26) and Dr. Nicole Beauchemin (ncxC26) for graciously providing cell lines used for our studies. Graphical illustration was created with Biorender. This work was delivered as part of the CANCAN Team supported by the Cancer Grand Challenges partnership funded by Cancer Research UK (CGCATF-2021/100034) and the National Cancer Institute (OT2CA278654). Other funding sources include those from the Ludwig Princeton Branch, and NIH grant CA163591 (to E.W.).

## AUTHOR CONTRIBUTIONS

Y.-Y.K designed and performed the experiments on the C26 model, analyzed data, and wrote the manuscript. Y.L. performed the food intake measurement. M.G.J. performed glucose flux analysis in the KP and KL model. M.A. assisted in glucose flux analysis in the KL model. G.J. and J.H. each assisted with mouse tissue collection and sample preparation for LC-MS. D.Y.L. contributed to flux data interpretation. T.J. contributed to flux data interpretation and revised the manuscript. M.D.G. supervised flux study in the KL model, contributed to flux data interpretation, and revised the manuscript. E.W. conceived and supervised flux study in the KP and KL models, contributed to flux data interpretation, and revised the manuscript. S.H. conceived and supervised the project and wrote the manuscript.

## DECLARATION OFINTERESTS

The authors declare no competing interests.

## SUPPLEMANTARY FIGURES

**Figure S1.**
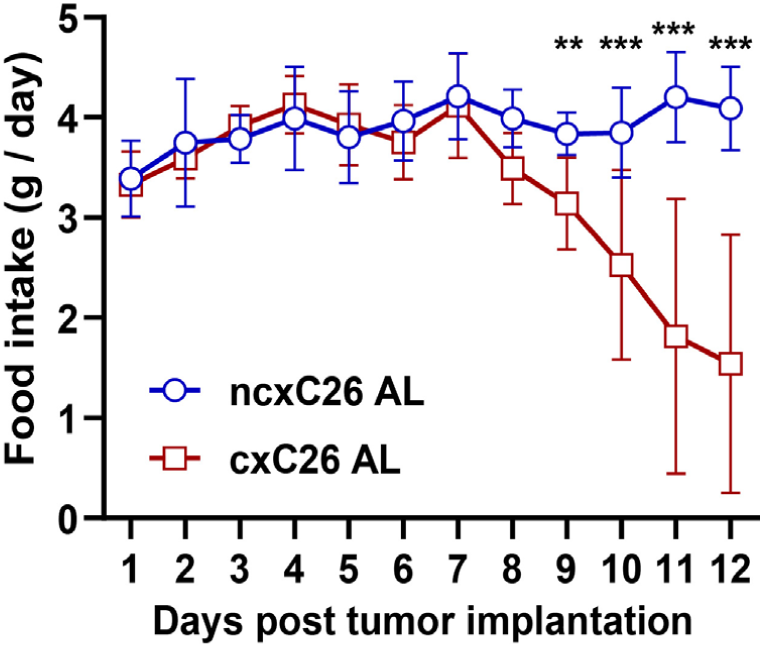
Food intake data of the C26 cancer cachexia model in this study. Daily Food intake was measured for individually housed cxC26 and ncxC26 mice ad libitum post tumor implantation. Any remaining food chunks and powder in the cage were collected and weighed to accurately calculate daily food intake. (n= 8) Data are shown as mean ± s.d. Significance of the differences: *P < 0.05, ** P < 0.01, *** P < 0.001 between groups two-tailed t-test.

**Figure S2.**
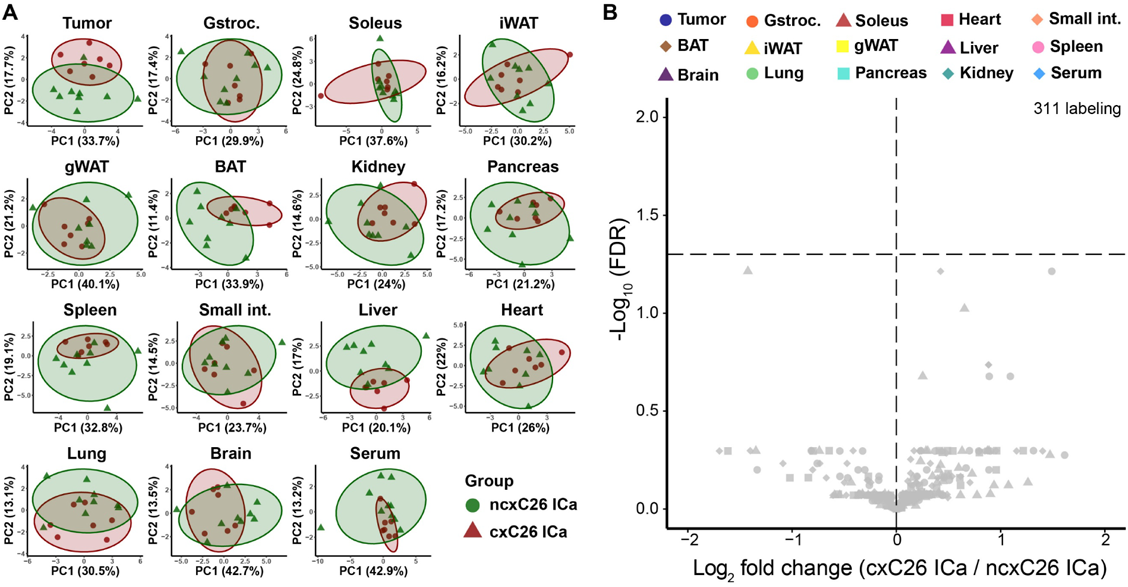
Labeling of metabolites in tissues of cachectic and non-cachectic mice under ^13^C-palmitate infusion. (A) Principal component analysis (PCA) of labeled metabolites in tissues of cxC26 and ncxC26 mice under ^13^C-palmitate infusion. (B) Volcano plot of differential labeling of tissue metabolites between cxC26 and ncxC26 mice under ^13^C-palmitate infusion. Statistical test for differences between cxC26 and ncxC26 groups was performed for each tissue, with the symbol (#) on a PCA plot indicating significance for that tissue (FDR < 0.01, MANOVA) and no symbol indicating no difference. All data was obtained for mice at day 12 post tumor implantation under isocaloric feeding (ICa). (n= 7-9)

**Figure S3.**
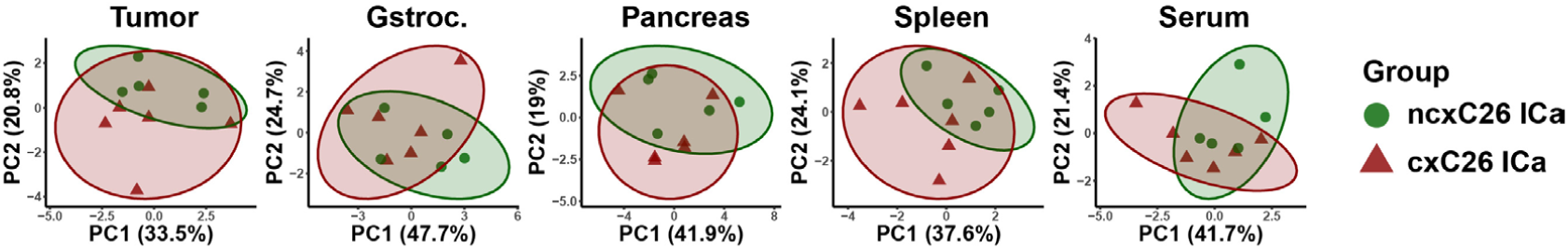
Labeling of metabolites in tissues of cachectic and non-cachectic mice under ^13^C-valine infusion. Principal component analysis (PCA) of labeled metabolites in tissues under ^13^C-valine infusion for cxC26 and ncxC26 mice under isocaloric feeding (ICa), at day 12 post tumor implantation. Only tissues with labeled metabolites detected are shown. (n= 5-6) Statistical test for differences between cxC26 and ncxC26 groups was performed for each tissue, with the symbol (#) on a PCA plot indicating significance for that tissue (FDR < 0.01, MANOVA) and no symbol indicating no difference.

**Figure S4.**
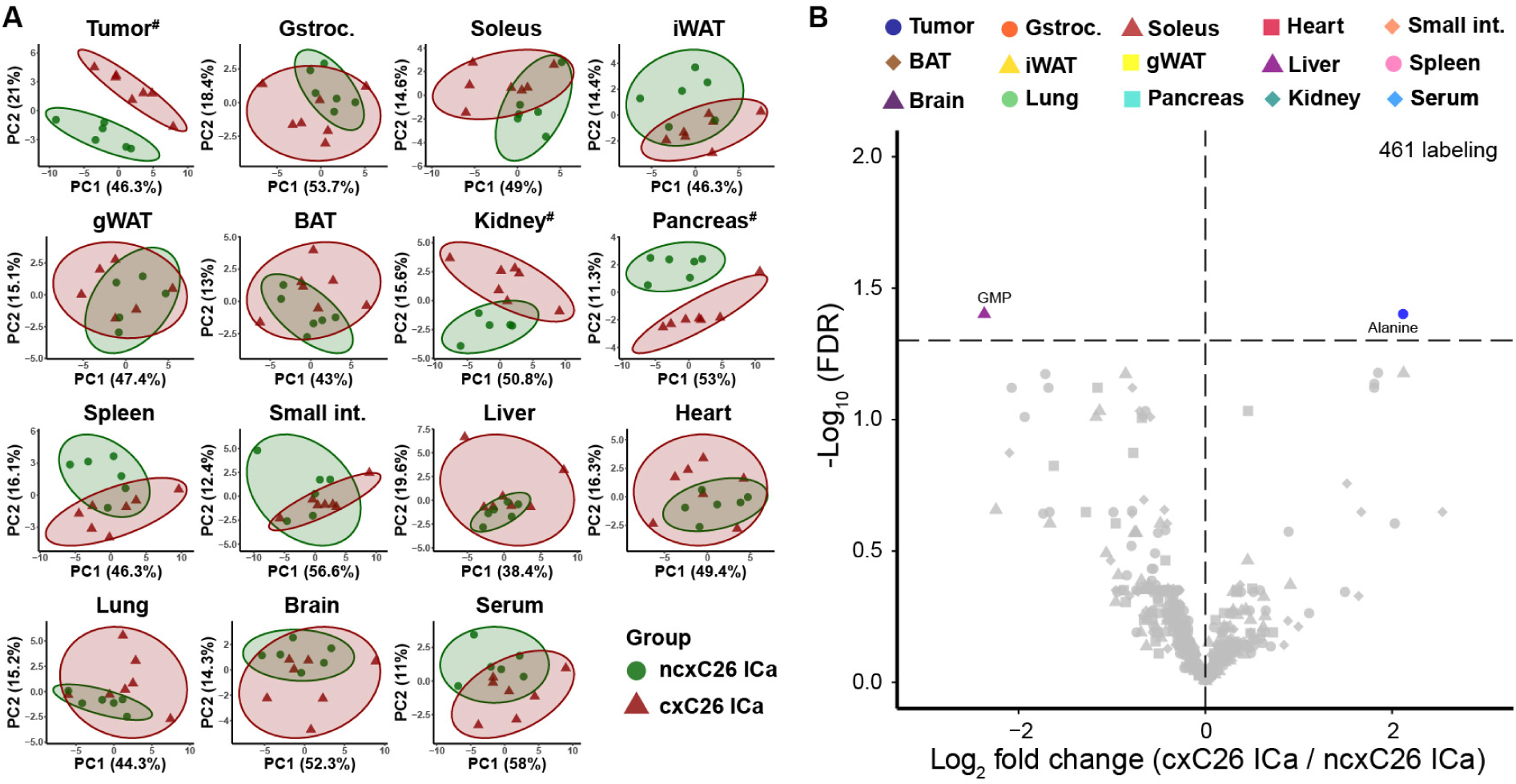
Labeling of metabolites in tissues of cachectic and non-cachectic mice under ^13^C-3-hydroxybutyrate. (A) Principal component analysis (PCA) of labeled metabolites in tissues of cxC26 and ncxC26 mice under ^13^C-3-hydroxybutyrate (3-HB) infusion. (B) Volcano plot of differential labeling of tissue metabolites between cxC26 and ncxC26 mice, under ^13^C-3-HB infusion. Statistical test for differences between cxC26 and ncxC26 groups was performed for each tissue, with the symbol (#) on a PCA plot indicating significance for that tissue (FDR < 0.01, MANOVA) and no symbol indicating no difference. All data was obtained for mice at day 12 post tumor implantation under isocaloric feeding (ICa). (n= 6-7)

**Figure S5.**
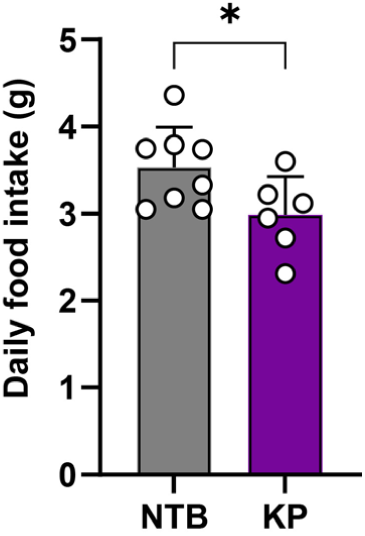
Food intake of the KP lung cancer cachexia model. Daily food intake was measured for cachectic KP mice and non-tumor bearing (NTB) mice when they developed cachexia (13 weeks post infection). Data are shown as mean ± s.d. Significance of the differences: *P < 0.05, between groups two-tailed t-test. (n= 8-9)

**Figure S6.**
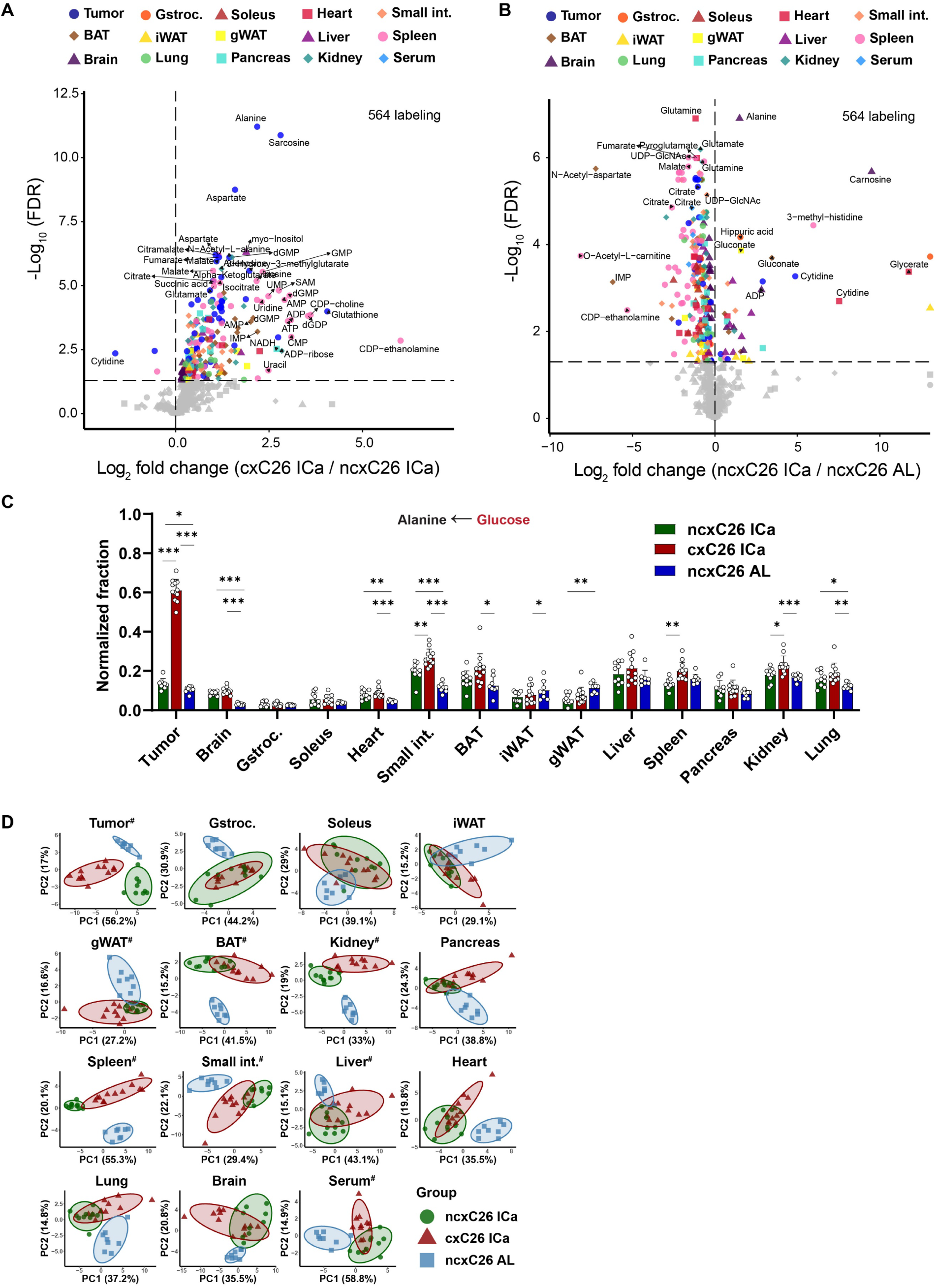
Labeling of tissue metabolites in cachectic and non-cachectic mice under ^13^C-glucose infusion. (A) Volcano plot of differential labeling of tissue metabolites between cxC26 and ncxC26 mice under isocaloric feeding (ICa), under ^13^C-glucose infusion. (B) Volcano plot of differential labeling of tissue metabolites between ncxC26 under isocaloric feeding (ICa) and ncxC26 ad libitum (AL), under ^13^C-glucose infusion. (C) Labeling of alanine in tissues under ^13^C-glucose infusion. no symbol: not significant, *FDR < 0.05, ** FDR < 0.01, ** FDR < 0.001 between groups by two-tailed t-test with multiple corrections. (D) Principal component analysis (PCA) of labeled metabolites under ^13^C-glucose infusion in tissues of cxC26 and ncxC26 mice. Statistical test for differences between cxC26 and ncxC26 groups was performed for each tissue, with the symbol (#) on a PCA plot indicating significance for that tissue (FDR < 0.01, MANOVA) and no symbol indicating no difference. All data was obtained for mice at day 12 post tumor implantation. Data are shown as mean ± s.d. (n= 8-11)

**Figure S7.**
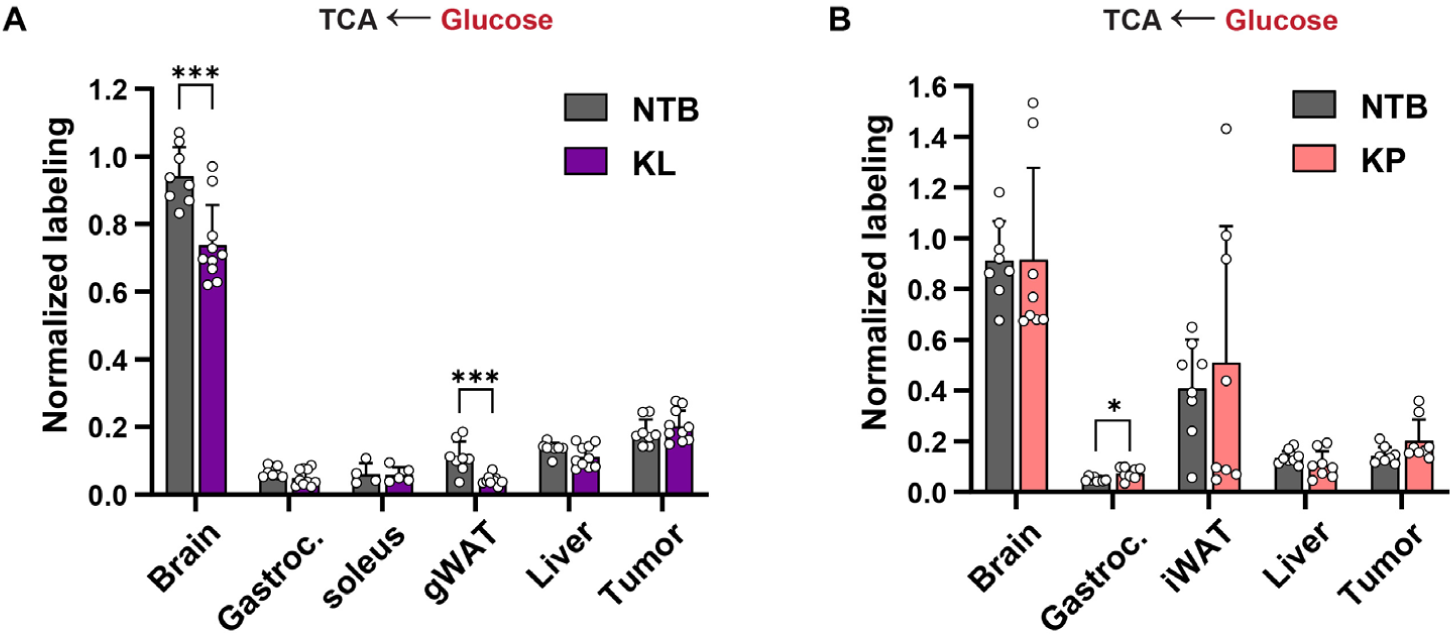
Labeling of tissue TCA cycle in KL and KP lung cancer cachexia models under ^13^C-glucose infusion. (A) Normalized labeling of malate in tissues of cachectic KL mice and non-tumor bearing (NTB) mice under ^13^C-glucose infusion. (n= 4-10) (B) Normalized labeling of malate in tissues of cachectic KP mice and NTB mice under ^13^C-glucose infusion. (n= 8) Data are shown as mean ± s.d. no symbol: not-significant, *P < 0.05, ** P < 0.01, *** P < 0.001 between groups by two-tailed t-test.

**Figure S8.**
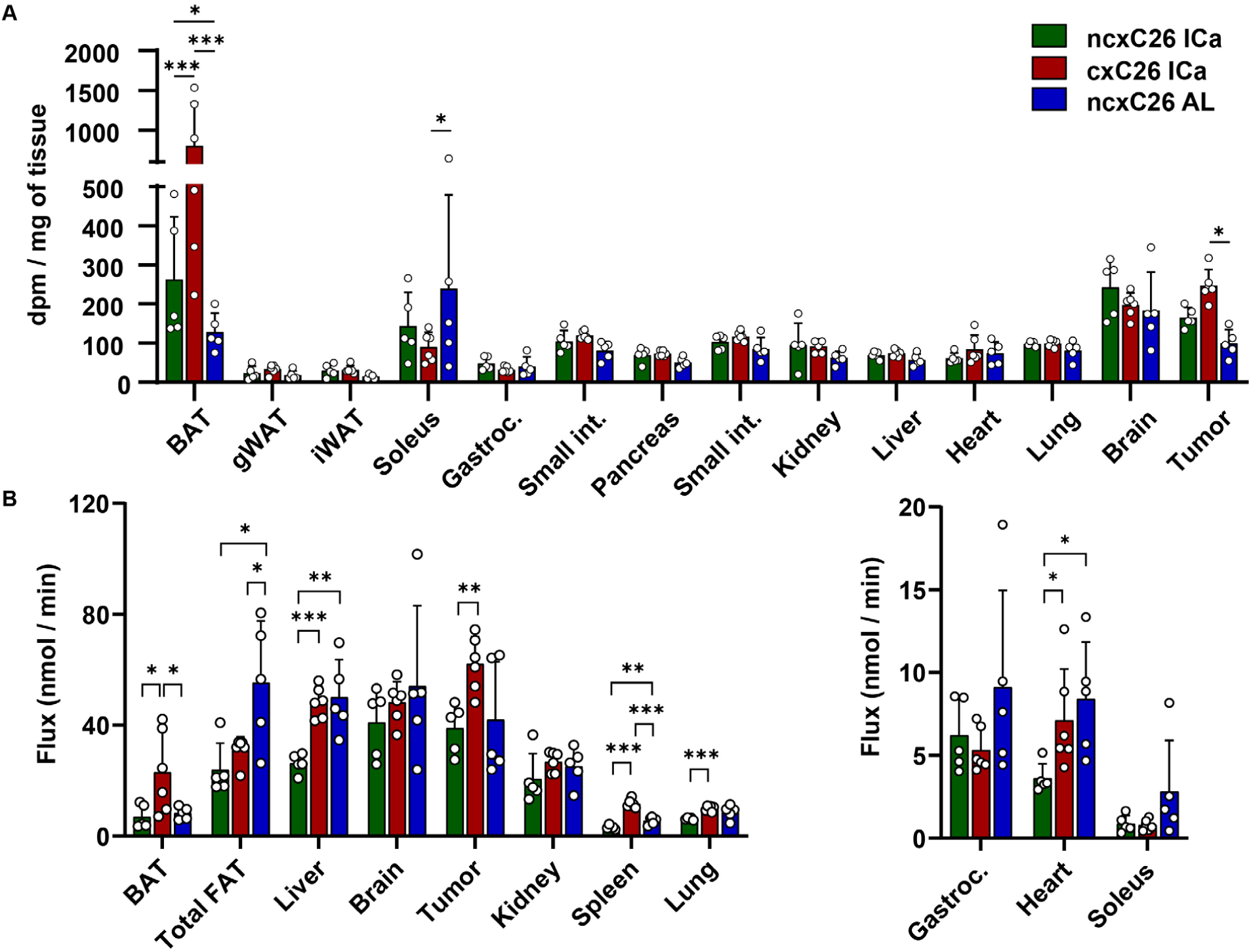
Tissue glucose uptake flux in cachectic and non-cachectic mice. (A) Uptake of ^14^C-2-Deoxy-D-glucose radioactivity by a milligram of tissues. (B) Calculated glucose uptake flux by tissues of cxC26 and ncxC26 under isocaloric feeding (ICa) and ad libitum (AL), at day 12 post tumor implantation. Data are shown as mean ± s.d. no symbol: not significant, *P < 0.05, ** P < 0.01, *** P < 0.001 between groups by two-tailed t-test. (n= 5-6)

**Figure S9.**
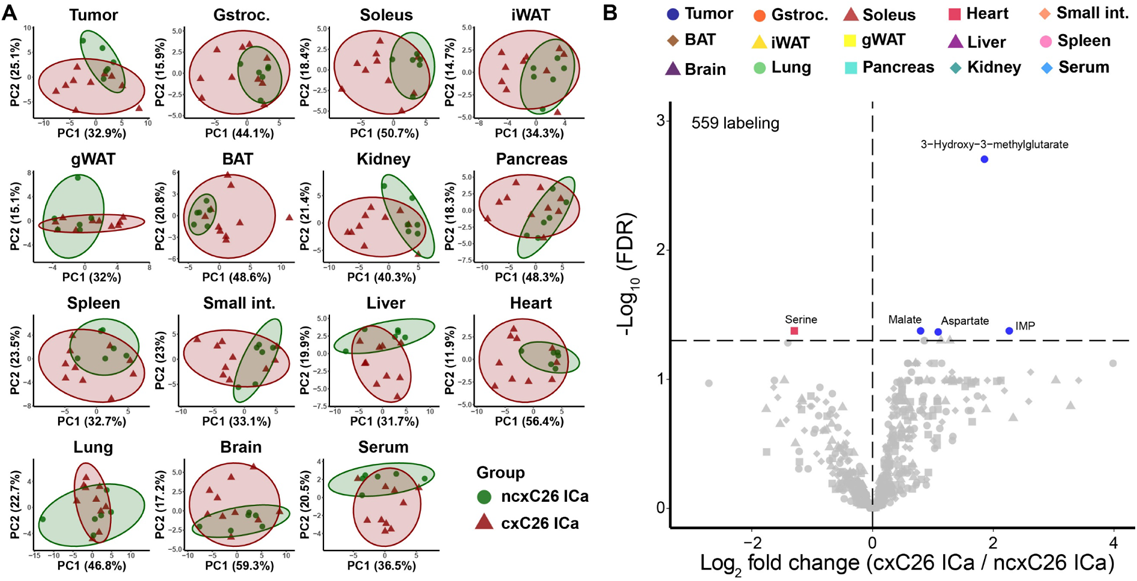
Labeling of tissue metabolites in cachectic and non-cachectic mice under ^13^C-lactate infusion. (A) Principal component analysis (PCA) of labeled metabolites in tissues of cxC26 and ncxC26 mice under ^13^C-lactate infusion. Statistical test for differences between cxC26 and ncxC26 groups was performed for each tissue, with the symbol (#) on a PCA plot indicating significance for that tissue (FDR < 0.01, MANOVA) and no symbol indicating no difference. (B) Volcano plot of differential labeling of metabolites in tissues of cxC26 and ncxC26 mice under ^13^C-lactate infusion. All data was obtained for mice at day 12 post tumor implantation under isocaloric feeding (ICa). Data are shown as mean ± s.d. (n= 6-13)

**Figure S10.**
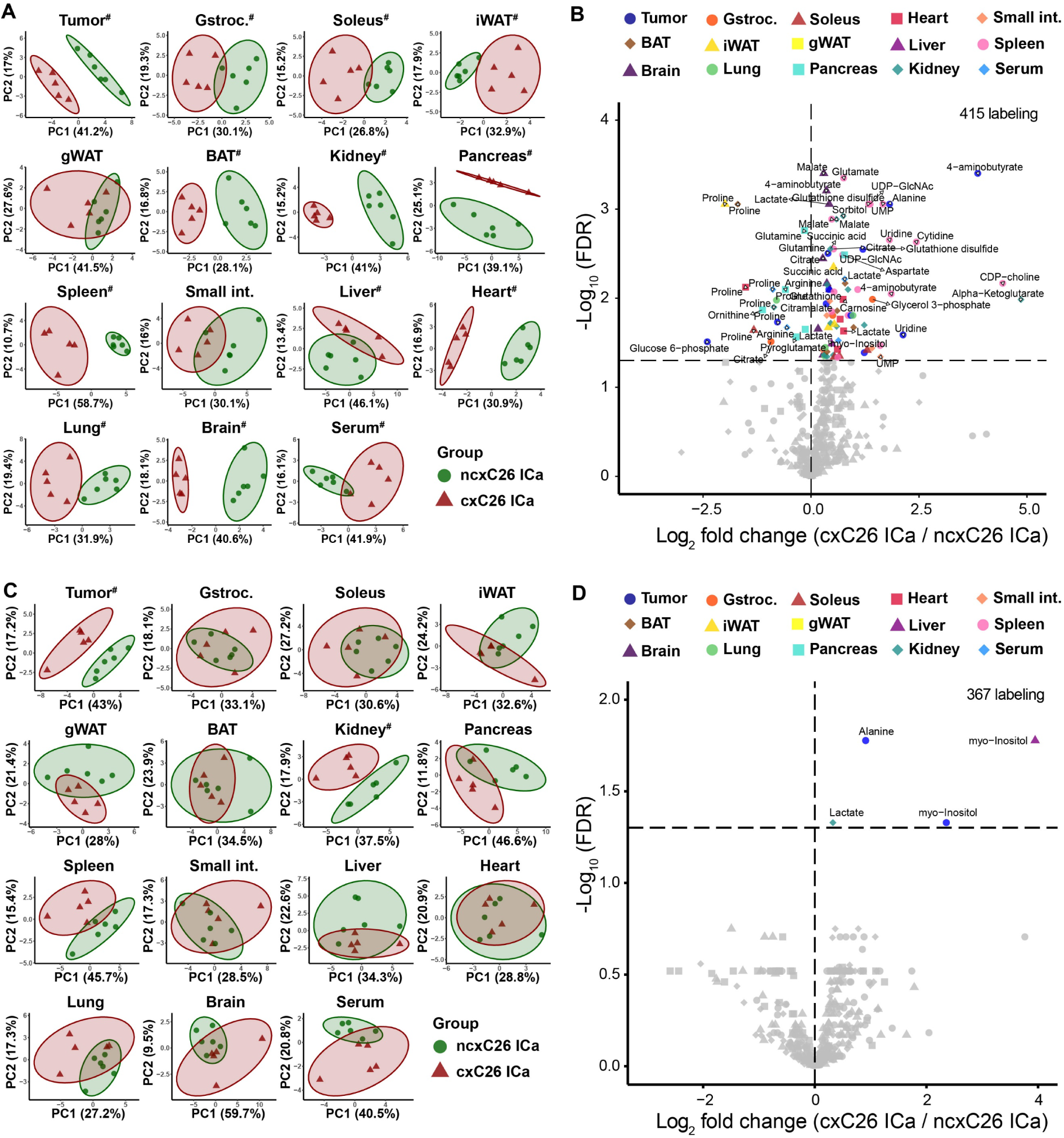
Labeling of metabolites in tissues of cachectic and non-cachectic mice under ^13^C-glutamine or ^13^C-alanine infusion. (A) Principal component analysis (PCA) of labeled metabolites in tissues of cxC26 and ncxC26 mice under ^13^C-glutamine infusion. (B) Volcano plot of differential labeling of metabolites under ^13^C-glutamine infusion in tissues of cxC26 and ncxC26 mice. (C) Principal component analysis (PCA) of labeled metabolites under ^13^C-alanine infusion in tissues of cxC26 and ncxC26 mice. (D) Volcano plot of differential labeling of metabolites under ^13^C-alanine infusion in tissues of cxC26 and ncxC26 mice. (A and C) Statistical test for differences between cxC26 and ncxC26 groups was performed for each tissue, with the symbol (#) on a PCA plot indicating significance for that tissue (FDR < 0.01, MANOVA) and no symbol indicating no difference. All data was obtained for mice at day 12 post tumor implantation under isocaloric feeding (ICa). (n= 5-6)

**Figure S11.**
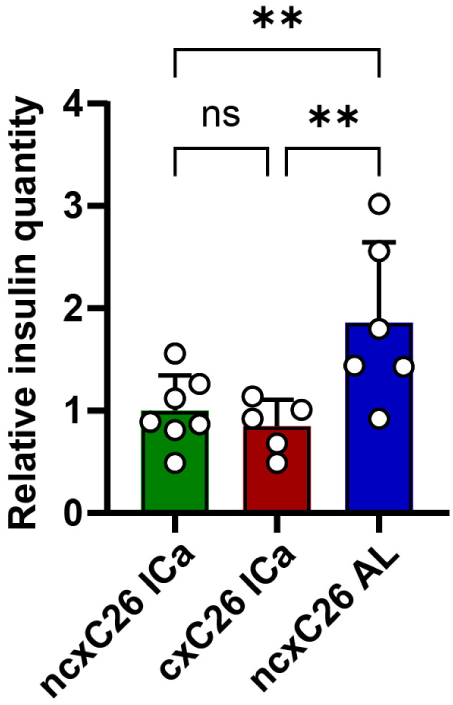
Circulating insulin was reduced to the same levels in cachectic mice and non-cachectic mice under isocaloric feeding. Relative circulating insulin levels in cxC26 and ncxC26 mice under isocaloric feeding (ICa), and ncxC26 mice ad libitum (AL), at day 12 post tumor implantation. Data are shown as mean ± s.d. Significance of the differences: ns: not significant, *P < 0.05, ** P < 0.01, *** P < 0.001, between groups by one-way ANOVA. (n= 5-7)

## Notes

### Competing Interest Statement

The authors have declared no competing interest.

